# Using deep convolutional neural networks to forecast spatial patterns of Amazonian deforestation

**DOI:** 10.1101/2021.12.14.472442

**Authors:** James Ball, Katerina Petrova, David A. Coomes, Seth Flaxman

## Abstract

1. Tropical forests are subject to diverse deforestation pressures but their conservation is essential to achieve global climate goals. Predicting the location of deforestation is challenging due to the complexity of the natural and human systems involved but accurate and timely forecasts could enable effective planning and on-the-ground enforcement practices to curb deforestation rates. New computer vision technologies based on deep learning can be applied to the increasing volume of Earth observation data to generate novel insights and make predictions with unprecedented accuracy.
2. Here, we demonstrate the ability of deep convolutional neural networks to learn spatiotemporal patterns of deforestation from a limited set of freely available global data layers, including multispectral satellite imagery, the Hansen maps of historic deforestation (2001-2020) and the ALOS JAXA digital surface model, to forecast future deforestation (2021). We designed four original deep learning model architectures, based on 2D Convolutional Neural Networks (2DCNN), 3D Convolutional Neural Networks (3DCNN), and Long Short-Term Memory (LSTM) Recurrent Neural Networks (RNN) to produce spatial maps that indicate the risk to each forested pixel (~30 m) in the landscape of becoming deforested within the next year. They were trained and tested on data from two ~80,000 km^2^ tropical forest regions in the Southern Peruvian Amazon.
3. We found that the networks could predict the likely location of future deforestation to a high degree of accuracy. Our best performing model – a 3DCNN – had the highest pixel-wise accuracy (80-90%) when validated on 2020 deforestation based 2014-2019 training. Visual examination of the forecasts indicated that the 3DCNN network could automatically discern the drivers of forest loss from the input data. For example, pixels around new access routes (e.g. roads) were assigned high risk whereas this was not the case for recent, concentrated natural loss events (e.g. remote landslides).
4. CNNs can harness limited time-series data to predict near-future deforestation patterns, an important step in using the growing volume of satellite remote sensing data to curb global deforestation. The modelling framework can be readily applied to any tropical forest location and used by governments and conservation organisations to prevent deforestation and plan protected areas.

## 1 Introduction

To achieve the pledge made by world leaders at COP26 to end deforestation by 2030, innovative approaches to forest protection are urgently required. Previous global efforts to curb tropical primary forest loss have so far failed to have a net positive impact (Potapov et al. 2017) despite countries and companies making substantial commitments (e.g. the New York Declaration of Forests in 2014). In the Amazon, deforestation is surging in some regions (Beuchle et al. 2021) and, unless it is halted in the next decade, there is a risk that the biome will undergo a critical transition to a savanna-like system with profound consequences for climate and biodiversity (Lovejoy and Nobre 2019). While the ultimate causes of forest loss need to be addressed – notably global demand for agricultural and wood products – interventions to tackle the proximate causes, such as illegal logging and settlement in protected areas, are also necessary. One such intervention that has been identified as being effective in reducing deforestation is “targeting protected areas to regions where forests face higher threat” (Busch and Ferretti-Gallon 2017). However, knowing how to optimally prioritise the allocation of limited resources to target diverse threats across often vast and difficult to access tropical forest regions is extremely challenging. To enable effective on-the-ground enforcement practices that curb deforestation rates, up-to-date information on the location, relative severity and likely evolution of threats is required.

Governmental and NGO commitments to make timely interventions have led to products that give near real time alerts on the location of ongoing or recent deforestation events (Matthew C. Hansen et al. 2016; Reiche et al. 2021), but access to these alerts has had little material benefit in terms of curbing deforestation (Moffette et al. 2021). One issue has been that responses based on near-real-time mapping can only ever be reactionary. As a result, there have been calls to build early warning systems that inform decision makers of the location of near-term deforestation risk and focus resources to high risk areas (e.g. (WWF 2020)). By developing innovative solutions that harness remote sensing data and emerging predictive technologies, interventions stand a greater chance of preventing illegal deforestation. In the longer term, cost effective conservation plans should not only consider the spatial distribution of conservation features (e.g. species / ecosystems, carbon stocks) and the costs related to their protection, but also account for how threats are likely to spread and evolve (Wilson et al. 2007; Boyd, Epanchin-Niell, and Siikamäki 2015). Predictive technologies can help to guide the spatial planning of protected areas.

Previous efforts to forecast locations of deforestation have had limited success often because they have been based on unreliable data layers and limited statistical or machine learning technologies. Deforestation is difficult to predict as it results from complex interactions within human-ecological systems but characteristic drivers have been identified (Geist and Lambin 2002) and spatially resolved geographical, economic, social and biological variables can indicate the likelihood of deforestation at a given location (Rosa, Ahmed, and Ewers 2014). Typically, within a statistical/machine learning framework, predictions are made by correlating several spatial predictor layers (e.g. distance to roads, agricultural land value) to the likelihood of deforestation based on the change in forest extent over a single time step of several years (see e.g. (Mayfield et al. 2017; Cushman et al. 2017; Saha et al. 2020) for intercomparisons of the efficacy of different statistical/machine learning frameworks). (Rosa, Ahmed, and Ewers 2014)’s review of such approaches concluded that they had a limited ability to accurately predict the future locations of deforestation, a key reason being that relevant, spatially explicit datasets on the drivers of deforestation are often incomplete or unreliable in tropical forest regions (especially those in the Global South). For example, proximity of roads is widely accepted to be a strong predictor of the location of future deforestation (Barber et al. 2014) but roads in tropical forests landscapes are often unofficial or not comprehensively mapped (Perz et al. 2007), and can be highly dynamic; appearing, expanding and changing course to access new resources (Ahmed, Ewers, and Smith 2014). Most studies have used static road maps as an input from which to predict deforestation which is inappropriate in many contexts, while others have used highly speculative methods to predict the development of roads in tropical forest landscapes (Ahmed, Ewers, and Smith 2014; Mena et al. 2017). With many of the decisions that lead to road development in tropical forest regions likely to remain hidden from public view, a fundamentally new approach to making predictions of the spread of deforestation is required to support effective local interventions. Analogous issues exist for other landscape features that can help to predict deforestation including the location/spread of human settlements and agricultural land. The statistical/machine learning frameworks used to date have also been limited in how they are able to represent local context. Without the ability to retain the spatial structure of the data, pixels are represented in isolation; what is happening close has been represented as simple averaged neighbourhood metrics. Contrasting approaches that identify threatened regions based on dynamic data (i.e. resolved over numerous time steps) have tended to focus simply on spatial and temporal trends in the intensity of deforestation (e.g. the emerging hot spot approach of (Harris et al. 2017)). While these approaches are easy to implement and update, they are naïve to the drivers of the observed forest loss (e.g. natural vs anthropogenic) and therefore unable to reliably predict whether a deforestation front is likely to spread which risks inefficient targeting of conservation resources to areas where true threats do not exist. A forecasting approach that could automatically learn and detect the contextual features of the landscape that signal deforestation risk from satellite imagery, and update its predictions accordingly, would bypass the limitations of both the statistical/machine learning and simple spatio-temporal trend based approaches.

By implementing modern computer vision approaches, it is feasible that an artificial neural network could automatically learn what features of a landscape are indicative of future deforestation from historic events and thereby avoid the need to rely on problematic data layers. Deforestation often exhibits striking spatio-temporal patterns when viewed from space (e.g. the classic fishbone pattern). The emergence of these patterns suggests some level of predictability while their characteristic spatial and spectral structures can give us clues as to the different drivers of deforestation. Over recent years, the quantity of freely available, high temporal and spatial resolution imagery has been increasing at a rate that has exceeded our ability to harness it for generating insights and supporting decision making. Deep convolutional neural networks (CNNs; a class of advanced machine learning models that maintains and work with spatial structures of data) that can be trained to automatically extract features of images have revolutionised the field of computer vision, allowing computers to reliably identify objects from images they are presented with (Voulodimos et al. 2018). They have led to radical advances in our ability to make predictions in countless fields from medical imaging (Yamashita et al. 2018) to solid-state materials science (Schmidt et al. 2019) and have recently been taken up by remote sensing researchers working with satellite data to generate novel insights (see (Zhu et al. 2017) for an overview), from remotely predicting levels of poverty (Jean et al. 2016), to quantifying the properties of terrestrial vegetation (Kattenborn et al.2021). 2D CNNs (designed for working with image data) (Kussul et al. 2017) and networks that incorporate an additional dimension (e.g. time) implicitly in their characterisation of data, including 3D CNNs (Y. Li, Zhang, and Shen 2017) and recurrent convolutional neural networks (ReCNNs) (Interdonato et al. 2019), have been shown to have state-of-the-art accuracy in detecting and classifying landcover change. Similarly, Long Short Term Memory (LSTM) based ConvRNN have been shown to improve on the performance of ReCNN for remote sensing classification tasks (Rußwurm and Körner 2018). These promising developments suggest that 2D CNNs, 3D CNNs and ConvRNNs architecture types, when trained on a large volume of spatially resolved historic deforestation data, could characterise and learn the features of the imagery associated with deforestation and be used to predict the future location of deforestation (something that has not previously been attempted). Critically, they are able to preserve and work with the spatial (and temporal) structures of the input datasets and therefore make predictions based on the local context of the scenes they are presented with.

This paper describes a range of original deep CNN model architectures that we designed to predict the risk of deforestation at a given individual pixel (30 m) when presented with a multi-layered scene (of data from freely available, global datasets) centred on that pixel, with the aim of automatically identifying areas threatened by deforestation. To test these networks, we trained and evaluated their performance on two regions in the Peruvian Amazon, one of the most biodiverse regions on the planet and exceptionally rich in endemic amphibians, birds, fishes, bats, and trees (Bass et al. 2010; Jenkins, Pimm, and Joppa 2013) but facing a range of threats including deforestation driven by copper and gold mining, logging, agriculture, cattle ranching and crude oil extraction (Finer et al. 2008; Piotrowski 2019). We trained and tested the models with cloud free satellite data for 2014-2020 and spatially resolved annual deforestation data for 2001-2020 (M. C. Hansen et al. 2013). The best performing model was used to forecast deforestation in these regions in 2021, predicting the risk of deforestation for every forested pixel in the landscape and classifying its likely future state accordingly. These spatial forecasts of deforestation risk could be a valuable asset for government agencies and NGOs who are attempting to make interventions that prevent future deforestation. The deep learning systems described herein (and available at **https://github.com/PatBall1/DeepForestcast**) can be readily reapplied and adapted to support conservation organisations in any tropical forest region.

## 2 Materials and Methods

### 2.1 Study regions

This study focused on two tropical forest regions in the south of Peru **Figure 1**). The sizes of the study regions were deemed suitable as they provided many millions of training points for our networks to learn from while allowing processing to remain within the computing resources available to us for reasonable experimentation. A preliminary intercomparison of a broad set of models (2D CNN, 3D CNN and LSTM ReCNN) focused on the Madre de Dios administrative department of Peru. Madre de Dios is almost entirely low-lying Amazonian rainforest (Olson et al., 2001). The Interoceanic Highway – a 2575-km road from the coast of Peru to Brazil that cost $2.8 billion to construct – was completed in 2011, allowing access to portions of Madre de Dios’s forest previously protected from human activity. Increased access has supported migration, typically from poorer Andean provinces, increasing pressure on the forest. Illegal gold mining is lucrative and prevalent in the region, seeding remote deforestation frontiers in otherwise pristine forest.

**Figure 1.**
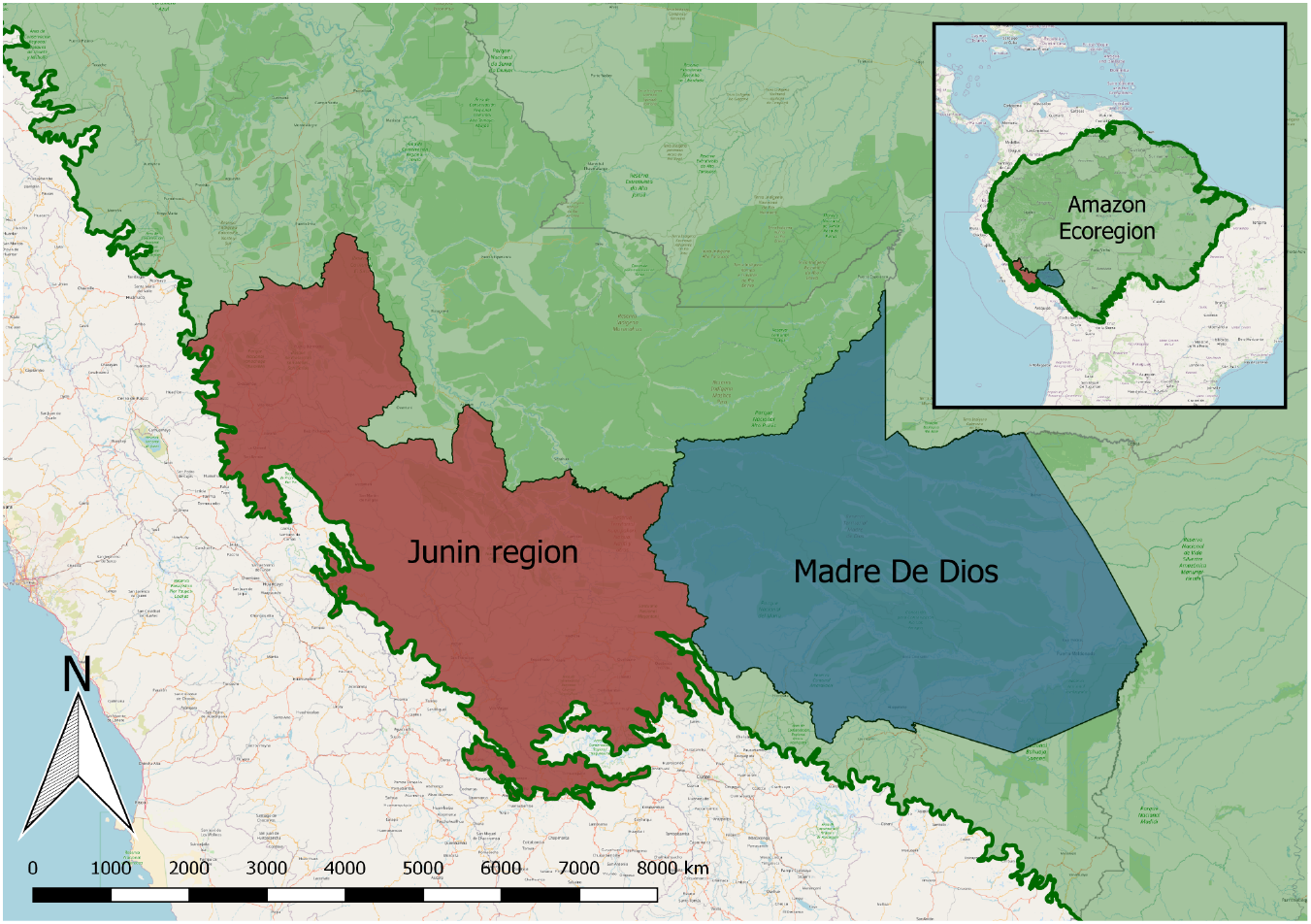
Southern Peruvian study regions. Madre de Dios is an administrative department of Peru and the Junin region is centred on the Junin department but includes other surrounding departments and is cropped by the limits of the Amazon ecoregion.

The most promising model architecture was selected, further developed, and applied to a second region. It centred on the Junín Department and included its surrounding departments and was cropped by the limit of the Amazon Ecoregion. This gave a similarly sized area but one with significantly greater topological heterogeneity to trail the networks on (see **Figure 1**).

### 2.2 Datasets

We limited ourselves to global, freely available datasets so the approach could be replicated for any forested location in the world. The code is open-source and freely available on GitHub (**https://github.com/PatBall1/DeepForestcast**) so anyone can implement the system to train a network and produce a forecast for their location of interest starting with just the spatial extent of the region (without required prior knowledge of machine learning). The code has been written so that a user can straightforwardly supply additional input data to train the models on and make predictions.

The forest cover labels used in this study were taken from the Global Forest Change dataset (Hansen et al., 2013). The dataset tracks the location of forest loss globally at 30m × 30m resolution and is collated on an annual basis (latest version: 2001-2020). The dataset includes a map of percentage tree canopy cover observed in the year 2000, a data mask map (describing areas of land or permanent water) and a cloud-free multi-spectral (Landsat) satellite image from the year 2000 which we included as predictor layers to help inform the network. The data is based on Landsat imagery and matches its resolution. Additional details on this dataset are given in **S2.** From the same source, pre-processed, cloud-free Landsat imagery of four spectral bands was available for every year from 2014 to 2020. The composite imagery takes the corrected spectral values for a pixel from the latest cloud free view of a pixel in a given year. A Digital Surface Model (DSM) provides a digital representation of the Earth’s surface. We included the Japan Aerospace Exploration Agency’s (JAXA) 30 m resolution ALOS Global DSM as a layer to give another dimension for the networks to learn and predict from, exploiting the features of the topology in the areas of interest. Historic deforestation labels (i.e. the year a pixel transitioned from forest to non-forest) were processed and included as four predictor layers to encode the proximity, in time and space, of recent deforestation and thereby assist the models in representing contagion effects. These one-hot encoded layers assigned pixels to a class based on how recently (from current time *t*) they became deforested: 0-1 years from *t*, 2-4, years from *t*, 5-8 years from *t* or more than 8 years from *t*. A pixel that does not register in any of these layers is considered free from deforestation since 2000. These data layers (see **Table 1**) were normalised to take values between 0 and 1 before being entered into the models. While additional layers could have been included to improve the accuracy of forecasts, we wanted to test the ability of the deep CNNs to extract useful features from a restricted and readily accessible set of datalayers.

**Table 1.**
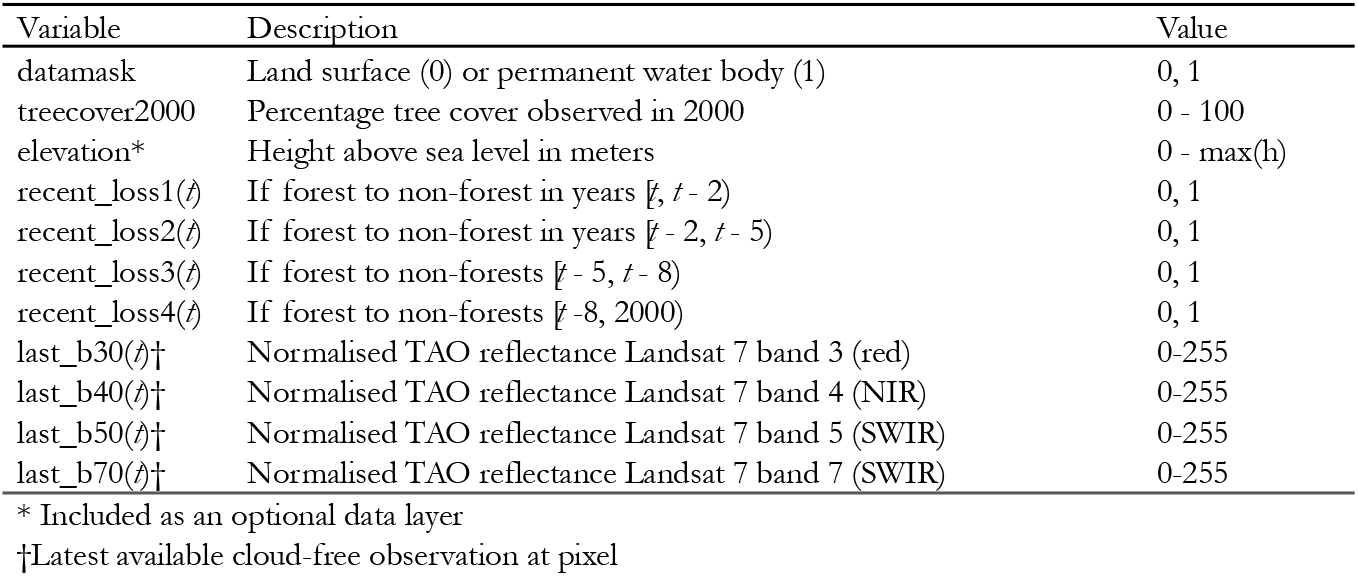
Predictor layers. The first three layers are static and the remaining ones change year-on-year.

| Variable | Description | Value |
| --- | --- | --- |
| datamask | Land surface (0) or permanent water body (1) | 0, 1 |
| treecover2000 | Percentage tree cover observed in 2000 | 0 - 100 |
| elevation* | Height above sea level in meters | 0 - max(h) |
| recent_loss1($t$) | If forest to non-forest in years $[t, t - 2)$ | 0, 1 |
| recent_loss2($t$) | If forest to non-forest in years $[t - 2, t - 5)$ | 0, 1 |
| recent_loss3($t$) | If forest to non-forests $[t - 5, t - 8)$ | 0, 1 |
| recent_loss4($t$) | If forest to non-forests $[t - 8, 2000)$ | 0, 1 |
| last_b30($t$)† | Normalised TAO reflectance Landsat 7 band 3 (red) | 0-255 |
| last_b40($t$)† | Normalised TAO reflectance Landsat 7 band 4 (NIR) | 0-255 |
| last_b50($t$)† | Normalised TAO reflectance Landsat 7 band 5 (SWIR) | 0-255 |
| last_b70($t$)† | Normalised TAO reflectance Landsat 7 band 7 (SWIR) | 0-255 |

\* Included as an optional data layer
† Latest available cloud-free observation at pixel

### 2.3 Modelling process

The aim was to train a classifier that, when presented with a scene centred on a single forested pixel, could predict whether the target central pixel will remain forested (0) or become deforested in the year ahead (1). The model training and testing used 2014 to 2020 data, and forecasts were produced for 2021 (a year for which data was not yet unavailable). This section contains a brief overview of the model training, validation and testing process. Full technical details are given in **S4** and corresponding scripts are available at [https://github.com/PatBall1/DeepForestcast].

At time *t* (years since 2000), the set of valid pixels(*J_t_*) were those that were in a forested state. These were the set of pixels on which training and prediction could be done. When a pixel transitioned from a forested state to a non-forest state, this change was taken to be permanent.

Specifically, a datapoint with a specific spatial location (the central pixel), *j*, and point in time, *t*, consisted of data from a **(2*r* + 1) × (2*r* + 1)** pixel scene, where ***r*** is the number of pixels the scene has in each spatial direction from the target central pixel, of the layers given in Table 1 (stacked as channels/bands). This allowed the network to observe what was happening in the neighbourhood of the pixel being considered (i.e. learn from and make decisions based on the field of view), which contrasts with typical statistical and machine learning approaches that have tended to use information on just the pixel in isolation or with averaged neighbourhood metrics that lose a lot of the available information. Each data point was associated with a non-forest (0) or forest (1) label from the year ahead (*t* + 1) (see **Figure 2**).

**Figure 2.**
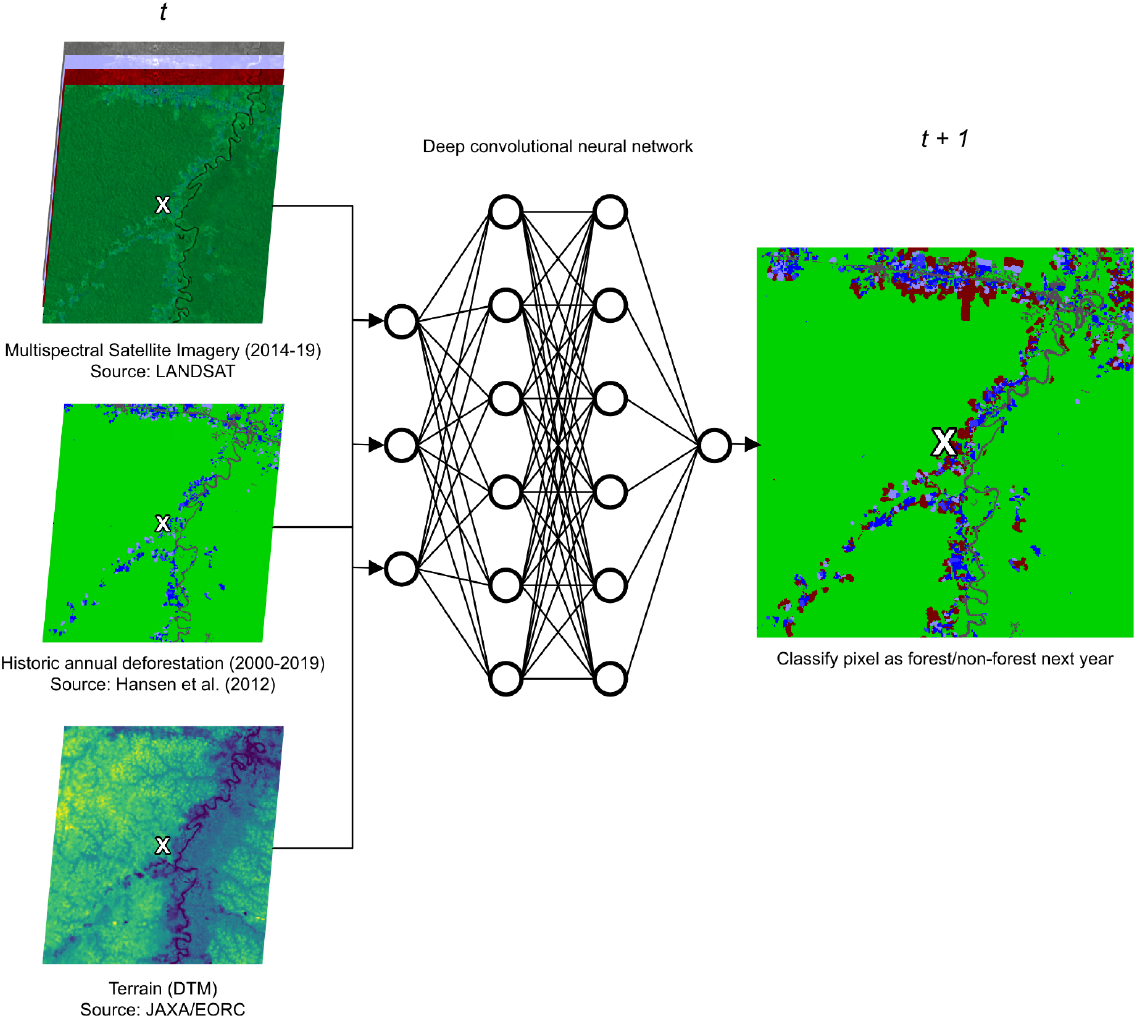
For a given forested pixel at the centre of a scene, the convolutional neural networks learned to predict the state of the pixel (forest/non-forest) in the year ahead based on multispectral satellite imagery, historic loss patterns and the terrain.

For each datapoint, two types of 3D tensors were inputted into the models. The first, ***S^j^***, contained the stacked static layers (see **Table 1**) and was of the shape ***S^j^* ∈ *R***^*s*×(2*r*+1)×(2*r*+1)^, where **s** is the number of static layers (2 or 3 in the current case). The second type, 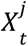, contained the stacked dynamic layers and was of the shape 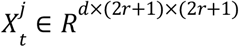 where *d* is the number of dynamic layers (5 in the current case). The label associated with the data point was defined as 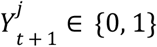, and took a value of 1 only if the central target pixel was labelled as non-forest in year*t* + 1. By allowing the models to learn the features (and interactions) of the data layers in the static and dynamic tensors that are associated with the year-ahead forest/non-forest labels, the model would then be able to make predictions of the year-ahead label 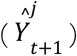 when presented with new input tensors. Output 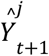 could vary continuously between 0 (no chance of forest loss) and 1 (certain transition to non-forest) with a threshold being applied to determine the binary prediction.

Less than 0.5 % of each study area transitioned from forest to non-forest in any given year which meant the number of valid pixels that remained as forest the next year (0) vastly outweighed the number that became deforested (1). This extreme class imbalance, if not addressed, would likely have led to very low predictive accuracy for the infrequent class (Ling and Sheng 2017). As suggested by (Buda, Maki, and Mazurowski 2018), we modified our training data by under-sampling the over-represented class so that there was at least one positive case (0 -> 1) to four negative cases (0 -> 0). Fortunately, even with this under sampling, there were several million valid data points available in each study region each year. All forest loss events were available for training or testing. The input data for each year, with associated year-ahead (*t* + 1) labels, was partitioned into training (80%) and test (20%) sets. The training set was further split into five even splits so that 5-fold cross validation could be performed to track model accuracy during the training process.

A preliminary, broad intercomparison of all model classes was first carried out on the Madre de Dios study region using 2014-2018 training and testing data. Using a grid search of hyperparameters, a broad set of models (different architectures and hyperparameters) was trained (with 5-fold cross validation) on 2014-2017 input data based on labels up to 2018. The accuracy of these models was tested on a (within training period) withheld test set (2014-2017 input data, 2018 labels). We used the test set AUC as a metric to compare the accuracy between the models.

Training deep convolutional neural networks is resource intensive so, from the initial intercomparison, the most promising model architecture was selected to be taken forward to be further refined and tested on the Junín region. Using Bayesian hyperparameter tuning (Snoek, Larochelle, and Adams 2012) and retraining on the 2014-2018 inputs (2019 labels), an optimised model was produced (the trails and performances achieved during the hyperparameter sweep is available to inspect at https://wandb.ai/patball/forecasting/sweeps/df5v36lz). The model was tested on the withheld 2014-2018 inputs (2019 labels) test set before being tested on the out-of-training-period test (2015-19 inputs; 2020 labels). It was important to test the models ability to predict the labels for a year which was outside of its training period to understand whether the model is transferable through time and therefore able to produce reliable forecasts. The optimised model (with parameters and weights retained) was then updated by continued training on 2015-2019 input data and 2020 labels. This final model was then used to produce deforestation forecasts across the entirety of each study area for 2021 (**available for inspection as a GeoTiff at https://doi.org/10.5061/dryad.hdr7sqvjz).** Using an earlier model version trained only on 2014-2017 (with 2018 labels), ‘forecasts’ for 2019 were also produced to compare against how deforestation truly evolved in 2019-2020 (**available for inspection as a GeoTiff at https://doi.org/10.5061/dryad.hdr7sqvjz**).

A fresh model, with an identical architecture and parameterization to the final model, was trained only on the Madre de Dios data. The accuracy of this model was evaluated first on Madre de Dios and then the Junin region to test spatial transferability of the model. This model was then used to forecast 2021 deforestation for Madre de Dios (**available for inspection as a GeoTiff at https://doi.org/10.5061/dryad.hdr7sqvjz)**. Another model with identical architecture and parameterization to the final model but sequentially trained on data on Madre de Dios data then Junin region data was assessed to help understand the effect of expanding the geographic diversity of training cases.

### 2.4 Model architectures

A variety of model architectures (MAs) were designed and implemented, and described in brief below. Full technical details and reasoning of choices of the network architectures can be found in**S3**. A deep neural network has a set of weights which define the strength of connections between the nodes and the predictions it makes from input data; these are adjusted as the network learns from exposure to training data. A single model architecture has a corresponding set of variable hyperparameters. Hyperparameters define aspects of how a model handles data and how it adjusts its weights to learn from the data. A unique set of hyperparameters as well as the architecture defines what we refer to as a *model*. By varying the hyperparameters and comparing the resultant model accuracies, the most effective set of hyperparameters can be identified (hyperparameter tuning; see above).

#### Model architecture 1: 2D CNN

The first and simplest MA we employed (MA1) was a type of 2D convolutional neural network (2D CNN). In a 2D CNN, filters slide (“convolve”) in two spatial dimensions across a 3D input tensor (with a spatial extent and a depth that corresponds to the number of bands) to extract spatial features that can be used by the network to make predictions. For any input into this model type a single dynamic tensor was combined with the static tensor so that the spatial features could be extracted. Potential temporal features of the data were not implicitly characterised by this type of network. Temporal change was captured by pairing the input data for one year (at time *t*) with the forest/non-forest labels from a year ahead (at *t* + 1). See **Fig S10** for a visual representation of the model architecture.

#### Model architecture 2: 3D CNN

The second MA (MA2) we employed was a type of 3D convolutional neural network (3D CNN). This type of network slides (“convolves”) filters in three dimensions – in this case two spatial and one temporal. This meant, for a given central target pixel, the series of dynamic tensors (see Section 2.5.3) could be stacked temporally along the channel axis to form a 4D input tensor and 3D convolutions used to characterise the spatio-temporal (spatial and temporal together) features of the input data. The static tensor was passed to a 2D convolutional branch. See **Fig S11** for a visual representation of this model architecture.

#### Model architectures 3 & 4: Convolutional Long Short Term Memory RNN

MAs 3 and 4 were also designed to implicitly handle the temporal characteristics of the input data. Instead of using 3D convolutions, they used a Convolutional Long Short Term Memory (ConvLSTM) based Recurrent Neural Network (RNN) type architecture. At a given time step, RNNs can *remember* and make use of information from previous time steps; they are designed to characterise and predict sequential data. LSTM is a specialised RNN design that allows for long time dependencies to be learnt. By combining this RNN architecture with spatial convolutional elements, a ConvLSTM can characterise and predict the sequential evolution of spatial patterns. MA3 had a single ConvLSTM cell whereas MA4 was “deeper” as it had a stack of ConvLSTM cells (see **S3.4** for details). These MAs take the same 4D input tensors as the 3D CNN and pass the static tensor to a 2D convolutional branch. These model architectures, while able to achieve comparable accuracies to the first two, were dropped after a broad intercomparison as their training could not be parallelised and so took far longer to achieve these accuracies. However, their model classes are available for experimentation from **https://github.com/PatBall1/DeepForestcast** as their implementation may become more effective as the amount of historic data and available computing resources increases.

### 2.5 Computation

We used the Adam optimizer with weighted cross entropy loss to train the models through stochastic gradient descent (Kingma and Ba 2014). The penalty for missing a loss event was set to be double that of incorrectly identifying a loss event at a pixel that remained forested as we felt it preferable for the network to be tuned to avoid missing the relatively rare loss events. To avoid overfitting, regularisation was implemented through a dropout layer (Srivastava et al. 2014) and early stopping criteria. The models were trained with repeated exposure to the full sampled training set (i.e. multiple *epochs*).

We used the PyTorch machine learning framework to construct, train and test the models (code available at https://github.com/PatBall1/DeepForestcast). We used multiple GPUs and CPUs running in parallel on a high performance computing cluster to train and test the models and make forecasts. Details on the computational resources used, including hardware and software, are given in **S5**. The trials conducted as part of the training process, including hyperparameters used, accuracies attainted and computing resources used is available at https://wandb.ai/patball/forecasting/.

## 3 Results

### 3.1 Model performances

#### Broad intercomparison (Madre de Dios)

The best models from each model class (i.e. selected after hyperparameter tuning) predicted 2018 Madre de Dios test set deforestation with similar accuracies (89%-90% when the test set was split 4:1 between pixel that remaining forested to deforested pixels as with the training data). Sensitivity and specificity were approximately equal across MA classes. The best performing MA after hyperparameter tuning was the 3D CNN (MA3; AUC = 0.944). This was followed by the deep ConvLSTM (MA4; AUC = 0.938) and the ConvLSTM (MA3; AUC=0.937). The 2D CNN architecture (MA1; AUC = 0.935) was the least accurate. This indicated that models that could implicitly handle and characterise the temporal with the spatial structure of the input data were better suited to predicting future deforestation. In other words, those models that could retain and work with the spatiotemporal patterns of the data (how the spatial patterns evolved) rather than just use spatial patterns tended to perform better. As well as being the best performing approach, MA2 also had a simpler architecture and was more suited to parallelisation than MA4 and MA3, and so was selected to be taken forward as the state-of-the-art MA.

The spatial dimensions of the input tensors had a strong impact on model performance as they determined the size of the scene that models were able to view. Model AUC plateaued around an input frame of ~ 35 × 35 pixels (~ 1 km × 1 km) and began to decrease for input frames larger than ~ 45 × 45 pixels (~ 1.35 km × 1.35 km). Between these sizes seemed to be the sweet spot where the networks could learn from local context while also remaining focused on the pixel of interest. Full details of the model performances, including hyperparameters and receiver operating characteristic (ROC) curves, are given in **S7**.

#### Focused development (Junín region)

Further tuning and extended training improved within-year accuracy the accuracy of the 3D CNN. Models that included elevation as an input predictor layer were no more accurate than those without. The information in this layer was either not helpful to the model in the form in which it was supplied, or the useful information that it did contain could readily be inferred from the other input layers. Following the principle of parsimony, this layer was dropped from the final training for the model to make the 2021 forecast.

As would be expected, there was a drop in model prediction accuracy from the within training period test set and the year ahead test set (see **Table 2**). This could indicate some change in the patterns of deforestation or a change in the nature of the input data (it is difficult to produce satellite data that is highly consistent across dates for a range of reasons including atmospheric conditions and cloud cover). The model that was trained on both regions sequentially was marginally better at predicting year ahead deforestation than the model that was trained on just the region to be predicted. Interestingly, the highest year ahead AUC for the Junin region was returned by the model trained exclusively on Madre de Dios data. This would indicate that the spatio-temporal patterns of forest loss between the two regions are somewhat similar and that the models are spatially transferable. It also suggests that data quality is more important than volume and spatial range of training data. The Junin region is far more mountainous than Madre de Dios and subject to very persistent cloud cover which could explain the apparent difference in data quality. The relatively poor performance of the Madre de Dios trained network on within year prediction for the Junin region suggests some amount of overfitting despite the implementation of regularisation techniques.

**Table 2.** A comparison of the within year and year ahead pixel wise classification accuracy of the 3D CNN model class. The 3D CNN model class was trained on different regions to test spatial transferability and the effect of increasing the volume of available training data. The reported accuracies are all for unseen test sets (2019 labels are within the training period and 2020 are outside). The ratio of 0 to 1 labels in the test set was 4:1.

| Model class | Training period (labels) | Training region | Prediction region | AUC 2019 | Accuracy 2019 | AUC 2020 | Accuracy 2020 | Sensitivity 2020 |
| --- | --- | --- | --- | --- | --- | --- | --- | --- |
| 3D CNN | 2014-2018 (2018-2019) | Junin | Junin | 0.989719 | 0.961253 | 0.838641 | 0.832001 | 0.580135 |
| 3D CNN | 2014-2018 (2018-2019) | Madre de Dios | Junin | 0.849717 | 0.829298 | 0.861616 | 0.832006 | 0.580015 |
| 3D CNN | 2014-2018 (2018-2019) | Madre de Dios; Junin | Junin | 0.996794 | 0.961630 | 0.849662 | 0.832352 | 0.580881 |
| 3D CNN | 2014-2018 (2018-2019) | Madre de Dios | Madre de Dios | 0.997370 | 0.984239 | 0.890692 | 0.886045 | 0.715114 |

### 3.2 Forecasts

The near-future forecasts exhibited some interesting spatial features. To illustrate the 3D CNN’s assessment of deforestation risk, we have focused on four recent forest loss events in the two study regions. The 2021 risk forecast was overlaid over Planet/NICFI biannual composite satellite imagery for 2020 (see **Figure 5**). **Figure 5a** shows a rapidly growing commercial agricultural area well serviced by a river and roads in Junín. The network signals a very high risk that agricultural clearance will continue to expand into the remaining forest areas. **Figure 5b** shows a newly laid access route connecting newly established agricultural settlements to a large river. The network highlights a risk of clustered conversion along parts of its route. **Figure 5c** shows a new, remote illegal gold mine in Madre de Dios. The network anticipates an immediate expansion of the mining operations into surrounding areas. **Figure 5d** shows a large recent forest loss event caused by a landslide. Despite the scale of the event, the network deems its surroundings as relatively low risk from further change of state when compared to the anthropogenic causes of loss. Indeed, a remote landslide is unlikely to result in a progressive loss of forest. The contrasting predictions (in combination with the high reported accuracy) suggest a level of discrimination (or *intelligence*) in the models evaluation of transition likelihood. While locations of likely future deforestation tended to be clustered around previous loss, the forecasts highlighted some moderate risk areas in scenes that had no recorded historic loss showing that forecasts were not entirely dependent on historic patterns of loss. Both forecasts are available for inspection in GeoTiff format at **https://doi.org/10.5061/dryad.hdr7sqvjz**.

**Figure 5.**
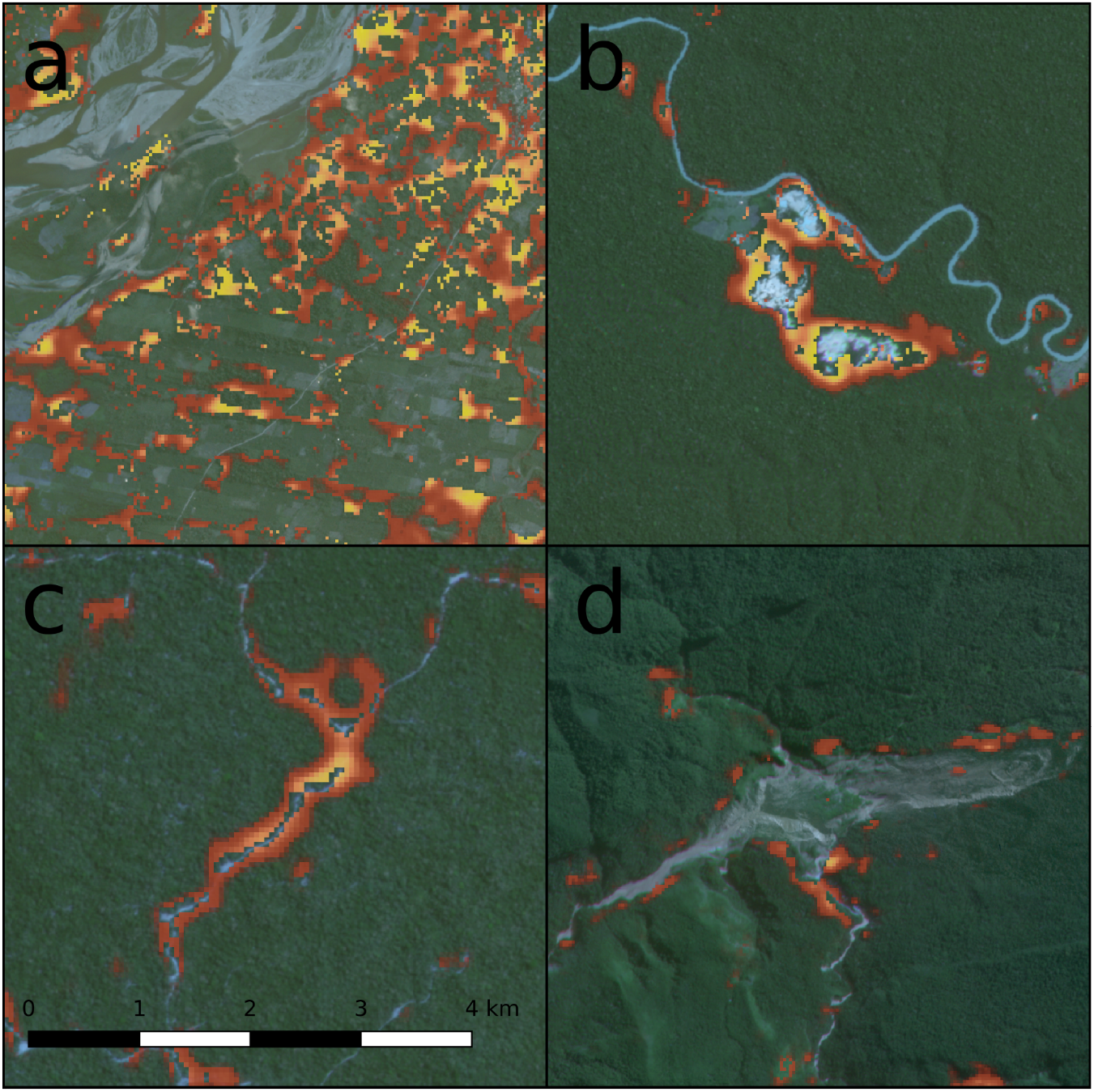
Examples from the Junín region and Madre de Dios 2021 deforestation risk forecasts where yellow indicates the areas most likely to transition to non-forest states fading to red as areas are less likely to transition. (a) shows a region of expanding commercial agriculture; (b) shows a new, remote indigenous settlement; (c) shows a new access route connecting a settlement to a river; (d) shows a recent remote landslide that led to a substantial clearing of forest. Source of satellite imagery: Planet/NICFI.

## 4 Discussion

For the first time, deep convolutional neural networks were used to forecast the spatial development of deforestation. From a limited set of freely available, global predictor layers, we predicted year-ahead deforestation at 30m resolution to a good degree of accuracy (80-90% pixelwise accuracy on a 4:1 negative to positive sampled test set) for two ~80,000 km^2^ regions in the Southern Peruvian Amazon while also demonstrating a degree of spatial and temporal transferability. By doing so, we have generated a new stream of data to support decision making and effective conservation interventions. Characterisation of the spatiotemporal structures of the data helped in the prediction of deforestation (relative to just the spatial structures as with 2D convolutions). The ability of these deep convolutional neural networks to automatically extract features from remote sensing data demonstrates the potential for deep learning computer vision methods to help us process remote sensing data to manage pressing risks in complex human-natural systems.

We stress that, while the reported accuracy is encouraging, there are reasons to take these figures with caution. The Global Forest Change (GFC) dataset provided all data labels for the training and testing of the forecasting system and so the accuracy specifically relates to how well the network predicts the future outputs of this classifier. The classification model used to generate the GFC dataset is a bagged decision tree that classifies filtered Landsat data and should not be considered a ground truth. Our models were likely to have approximated and incorporated behaviours of this classifier and accuracy may be inflated (relative to on-the-ground change) by learned correlations with this model. For example, there may have been cases where the network learned to anticipate a classification that the Global Forest Change classifier makes at a later date based on shifts in the spectral signal. Conversely, our reported accuracy may be conservative relative to the ground-truth; the deep neural network architecture would theoretically be able to pick up more subtle spectral signals than the less complex bagged decision tree.

The apparent ability of the networks to automatically extract landscape features to discern different drivers of forest loss (natural vs anthropogenic) and infer whether loss was likely to continue from a source was encouraging. Previous attempts at forecasting deforestation have often had limited predictive capacity due to a lack of reliable input data on landscape features such as roads. From satellite data and other complementary spatial datasets, our model appeared able to discriminate between a one-off forest loss event (e.g. landslide) and deforestation frontiers that are likely to progress over time (e.g. a newly built road, an illegal gold mine). This suggests that the networks were able to automatically extract those features of the landscape that indicate risk of deforestation. By focusing the training process more on contentious cases (i.e. those cases that sit on the threshold between the two classification states) it may be possible to further boost the accuracy of the network. The optimal window width to view the scene was 1 km - 1.35km. For larger windows, the marginal benefit from the additional information provided less benefit than the cost of handling this information. In other words, there was a point at which adding more further away pixels to a scene began to distract the network from the area of focus.

Deep neural networks are considered ‘black box’ models so it is difficult to say how the networks were making their decisions and to what degree they were recognising different potential drivers of loss. There has been work on interpreting deep CNNs to explore their internal reasoning (X. Li et al. 2021). By applying available interpretation algorithms it would be possible to identify which parts of a scene are triggering the networks’ decision. It may even be possible to generate groups of scenes (e.g. grouped by driver) that typically signal forest loss (see e.g. Activation Atlas (Carter et al. 2019)). Automatic classification of deforestation drivers through the application of machine learning to remote sensing has been demonstrated at coarse scales (10 km) with simple decision tree models applied to derived remote sensing products (Curtis et al. 2018). Given the relative sophistication of the CNN technology available, it is entirely feasible that a CNN could be trained to classify drivers directly from satellite imagery at high resolution (~10 m), particularly as the amount of freely available, high resolution imagery is rapidly increasing (e.g. Planet NICFI, Sentinel-2). However, a comprehensive training set of labelled deforestation drivers at this resolution does not exist and would require a considerable investment to produce. Integrating higher resolution into the forecasting system may help it to discern drivers more precisely but, just as there was an optimal window size, there may be an optimal resolution of imagery beyond which the additional information no longer adds value either due to the diminishing relevance of the information or computational/technical constraints.

The forecasting approach and outputs presented here were designed so as to be flexible and useful to a range of potential users. All pixels in the landscape can be ranked according to the likelihood of loss allowing for spatial conservation prioritisation to be performed. A government agency may have the resources available to police a certain proportion of the area under the jurisdiction and the generated forecasts would allow them to monitor the most threatened areas. The forecasts could also direct interventions to emerging frontiers of loss in otherwise isolated and intact landscapes; these are particularly problematic in terms of ecological impact and difficulty of prevention. On the other hand, when planning a permanent protected area, it may be preferable to write off the areas that are most imminently threatened but address areas that may undergo transition if left unprotected (i.e. address the medium risk areas). In either case, it is beneficial to have an understanding of how threats will evolve. This is also true in the rapidly growing field of forest carbon accreditation where short term forecasts could provide the counterfactuals against which the ‘additionality’ of projects could be measured. This does raise the issue that a forecast can fundamentally change the behaviour of the system that it is trying to predict and indeed this would be the case if interventions based on the forecasting system presented here were widely implemented. Were this the case, it would be possible to encode the interventions in the system as an additional spatial data layer and measure the impact of the interventions based on the relative change in likelihood of deforestation within/around the region of intervention (based on updated data and retraining of the models).

Any machine learning approach is necessarily limited by the fact it is based on historic data and the assumption that observed trends will continue, and therefore does not implicitly account for regime changes in the system. This is particularly relevant in this work, as political changes are known to have substantial impacts on Amazonian deforestation (Pereira et al. 2020). A changing climate, international economic conditions and advances in technology also challenge the assumption of stationarity. However, as the system outputs a continuous metric of risk across all pixels, by varying the threshold at which the binary classification is made it is possible to adjust the overall allocation of forest loss. It would be possible to couple the direct spatial evaluation of risk from the CNN system (that can account for sudden changes at the landscape scale) to a model of overall forest loss based on external (e.g. economic) drivers to generate forecasts that reflect global trends.

So far, it has only been feasible to train on satellite imagery as far back as 2014 but training on a deeper time series would allow the networks to learn from a greater range of contexts and would likely produce more accurate forecasts. However, years in the more distant past may not be as representative as recent years with respect to deforestation patterns so it is unclear as to whether extending the training period would significantly boost performance. It would also be of value to understand whether using higher resolution satellite imagery (e.g. Sentinel-2, Planet NICFI) would allow the network to pick up on smaller scale features of the landscape and predict more accurately but, as mentioned above, this would involve a trade-off.

The networks were designed to produce just +1 year forecasts. Due to the inherent stochasticity in the system long-term forecasts (5+ years) would not be feasible to any degree of accuracy. However, it would be valuable to test the effect of extending the forecasting horizon on accuracy to up to +3 years.

The model architectures were necessarily designed from quite fundamental components, bespoke for a very particular task. However, this approach foregoes some of the innovations that have been included in other state-of-the-art approaches (e.g. *attention* as in CoAtNet (Dai et al. 2021), the best model on the ImageNet classification benchmark). Further research could be done on integrating architectural components that have been developed by leading AI teams. Indeed, we hope that this paper highlights the potential of deep CNN techniques for a task of great ecological and social value and that others in the field may continue to improve the efficacy of the approach by continued technological innovation. By further improving the sophistication of the network it may be possible to achieve the improvements suggested above.

In its current form, the forecasting approach should be applied to and tested on other forest regions, both tropical and temperate. It would be helpful to test how performance differs in regions with different ecological/social contexts and drivers of deforestation. Additional predictor layers can be readily integrated so that researchers can adapt the system based on their goals.

Governmental and non-governmental organisations can immediately use the tools presented here to help understand how the landscapes in which they operate will change into the future, and support their engagements with local communities to mitigate carbon emissions. Additionally, local and national governments could incorporate the system into their protocols for managing and responding to deforestation risk. Beyond this, the effectiveness of deep CNNs in the current context suggests that they could also be applied to other remote sensing tasks in the fields of forest ecology and conservation such as fire detection and forecasting and forest restoration.

## 5 Acknowledgements

Thanks to Mark Burgman for initiating the collaboration between the authors. Thanks to Cool Earth for their early support of the research. James Ball is supported by the Natural Environment Research Council. Seth Flaxman is supported by the Engineering and Physical Sciences Research Council.

## 7 Authors’ contributions statement

JGCB, SF and KP conceived of the ideas and designed methodology presented in the study. KP, JGCB and SF coded the models and routines. JGCB led the writing of the manuscript. SF, DAC and KP were involved in writing and reviewing the manuscript.

## 8 Data Availability

All code is freely available on Github. Records of model runs and hyperparameter tuning is available at https://wandb.ai/patball/. All input data is freely available online. Processed, model ready inputs are available on **Dryad [https://doi.org/10.5061/dryad.hdr7sqvjz]**. Model weights and specifications are available on **Dryad [https://doi.org/10.5061/dryad.hdr7sqvjz]**. Using these with the model ready inputs, code, and virtual environment specification (available on Github), all model outputs are reproducible. The code is designed so that a user can run the full training, testing and forecasting process including the initial downloading and processing of input data.

