## Supplementary Materials for "Using deep convolutional neural networks to forecast spatial patterns of Amazonian deforestation"

### Spatially Forecasting Deforestation with Deep Neural Networks: Supplementary Materials

*Affiliations:*
[1] Department of Plant Sciences, University of Cambridge
[2] Department of Mathematics, Imperial College London
[3] Deptartment of Mathematics, University of Oxford

November 2021

**References** **43**

### S1 Delimitation of study areas

#### S1.1 Madre de Dios

The boundary of Madre de Dios was taken from a shapefile of Peru's administrative Departments[1].

#### S1.2 Junin region

The study region was defined by the following steps: Starting layers were:
1) Shapefile of Peru's administrative Departments
2) WWF map of the entire Amazon Ecoregion

Junin, and the five surrounding departments, Pasco, Apurimac, Ayachcho, Cusco and Huancavelica, were selected and merged (dissolved) into one area. Then, the intersection of the dissolved departments and the Amazon Ecoregion was calculated. Some small islands detached from the main area were removed. To allow for predictions close to the edge of the region, a second region was also defined by adding an additional buffer of 0.09 degrees and the removing any internal islands that had not been included.

[1] https://data.humdata.org/dataset/limites-de-peru

### S2 Datasets and feature extraction

#### S2.1 Global Forest Change dataset and Landsat imagery

The Global Forest Change dataset (Hansen et al., 2013) gives areas (at approximately 30×30 meter resolution) that have been detected from Landsat imagery as undergoing forest loss. The first release of the global map of forest cover loss, was made in 2013 and contains a map of forest extent (in 2000), annual loss (from 2000 to 2012), and overall gain (from 2000 to 2012). Since then, the data set has been updated annually with the same methodologies being used as in 2013. The current Version 1.8, includes forest loss from 2000 through to 2020. The features of the data is given in table S1.

"Tree cover" is defined as "all vegetation greater than 5 meters in height, and may take the form of natural forests or plantations across a range of canopy densities". Forest loss is defined as "the disturbance or complete removal of tree cover canopy (below 30% tree canopy cover)". The forest loss detection does not differentiate between permanent tree cover loss or temporary loss from which the forest will recover. It also does not determine whether the cause of the loss is natural or human induced. The dataset comes with a layer of percentage tree canopy cover observed in the year 2000, a data mask map (describing areas of land or permanent water) and a cloud-free multi-spectral (Landsat) satellite image from the year 2000. Additionally, each annual release comes with the most recent available cloud-free Landsat image composite from that year (the "last" layer) and updated layer an annual forest loss ("lossyear").

The latest data was extracted for the Madre de Dios area and the Junin area described above. Consistent cloud-free Landsat imagery was available from 2014. Annual forest loss events were available from 2001 to 2020.

**Table S1:** Layers of the Global Forest Change dataset

| Variable | Description | Value |
|---|---|---|
| treecover2000 | Percentage of tree cover in the pixel observed in 2000. | 0 - 100 |
| gain | One if gain happens during the period: 2000 - 2012, zero otherwise. | 0 or 1 |
| lossyear | The year when loss was detected, one-indexed from year 2001, or zero if no loss occurred. | 0 - 18 |
| datamask | No data (0), mapped land surface (1), and permanent water bodies (2). | 0,1 or 2 |
| first_b30 | The Landsat 7 red band built from the first cloud free pixels in 2000. | 0 - 255 |
| first_b40 | The Landsat 7 near infrared band built from the first valid pixels in 2000. | 0 - 255 |
| first_b50 | The first Landsat 7 short wave infrared band built from the first valid pixels in 2000. | 0 - 255 |
| first_b70 | The second Landsat 7 short wave infrared band built from the first valid pixels in 2000. | 0 - 255 |
| last_b30_2013 (/2014/..) | The Landsat 7 red band built from the latest valid pixels in 2013 (/2014...). | 0 - 255 |
| last_b40_2013 (/2014...) | The Landsat 7 near infrared band built from the latest valid pixels in 2013(/2014...). | 0 - 255 |
| last_b50_2013 (/2014/..) | The first Landsat 7 short wave infrared band built from the latest valid pixels 2013 (/2014...). | 0 - 255 |
| last_b70_2013 (/2014/..) | The second Landsat 7 short wave infrared band built from the latest valid pixels 2013 (/2014...). | 0 - 255 |

Since the utility of optical satellite imagery is highly reliant on cloud cover (which can be persistent across the Andean Amazon), some late-in the year losses may only detected in the following year. The assignment of deforestation events to a particular year should therefore be treated with some caution.

Additionally, two different algorithms were used to generate the the measurements of tree cover loss - one for 2001-2010 and another for 2011-2018. The new algorithm is more sensitive to small-scale agricultural, fire-caused, or other forest losses. Goldman, Weisse (2019) noted that they observe large spike in the tree cover loss in the years 2016 and 2017 globally, and that the causes for that are determined to be fires. While this can be considered as an anomaly in their dataset, they also suggest that other pre-2011 fire-related losses may not detected by their initial algorithm. The scientists involved in this project are working to back-cast the new algorithm to generate one consistent time series of forest loss events.

#### S2.2 Digital surface model

The second set of models for the Junin/Ashaninka area were further developed by the inclusion of an elevation predictor layer to allow the models to learn from features of the topology.

A Digital Elevation Model (DEM) or Digital Surface Model provides a digital representation of the Earth's surface. In the We included the Japan Aerospace Exploration Agency's (JAXA) 30-m resolution ALOS Global DSM as a layer for the models. Including a DSM as a layer gives another dimension to the data and allows the deep networks to learn and predict from features of the topology of the areas of interest.

A review of the available DEM/DSM sources was performed. Alternatives included NASA's SRTM DSM, the MERIT DEM and the TanDEM-X DEM. The later two were excluded as their resolution was too low. SRTM is said to struggle in sloping regions with foreshortening, layover and shadow. JAXA's ALOS Global DSM was judged to be the most precise and suitable to complement the Global Forest Change and satellite data described above.

#### S2.3 Feature extraction

Global Forest Change dataset (Hansen et al., 2013) is divided into 10x10 degree tiles, each of which comes with six raster files per tile: treecover, gain, data mask, loss year, first and last (see Table S1). All files contain unsigned 8-bit values and have a spatial resolution of 1 arc-second per pixel, which correspond to approximately 30 meters per pixel around the equator. After 2013 loss year and last files were updated annually. The last 2020 loss year file assign an integer value 0-20 to each pixel. 1-20 corresponds to the year (2001-2020) at which a forest to non-forest event was observed at this location. 0 is assigned if no change is detected in the period 2001-2020. The dataset is encoded such that once a pixel is assigned as deforested, it does not go back to forested at any time in the future. We collected the following ten tif files: treecover, gain, datamask, all "last" files from 2014 to 2020 and the most recent, 2020, loss year file. Since we wish our models to be able to predict the label of each pixel of the regions by analyzing an image, or time series of images, that captures its local region, we also included pixels lying in a buffer area of 0.09 degree (or approximately 10km) in our dataset. This allowed us to extract features from images that cover area up to 10km away from within region pixel.

From the processed dataset we then assigned 7-8 predictor values to each pixel, 2 stationary and 5 that vary each of the years 2014-2020. Table S2 provides explanation for each of them. Each pixel also has a corresponding lossyear value $\in 0, 1, 2, 3, 4, ...20$, where 1-20 indicate the year at which it was marked as deforested or 0 if it did not experience deforestation up to year 2020.

**Table S2:** Predictors

| Variable | Description | Value |
|---|---|---|
| datamask | Mapped land surface (0), and permanent water bodies (1). | 0, 1 |
| treecover2000 | Percentage of tree cover in the pixel observed in 2000. | 0 - 100 |
| elevation* | Height above sea level in meters | 0 - max(h) |
| recent loss1(t) | If pixel transitioned from forest to non-forest in years [t, t - 2) | 0, 1 |
| recent loss2(t) | If pixel transitioned from forest to non-forest in years [t-2, t - 5) | 0, 1 |
| recent loss3(t) | If pixel transitioned from forest to non-forest in years [t-2, t - 5) | 0, 1 |
| recent loss4(t) | If pixel transitioned from forest to non-forest in years [t-2, t - 5) | 0, 1 |
| last_b30(t)† | Normalised TAO reflectance Landsat 7 band 3 (red) from the latest valid pixels in year t | 0 - 255 |
| last_b40(t)† | Normalised TAO reflectance Landsat 7 band 4 (NIR) from the latest valid pixels in year t | 0 - 255 |
| last_b50(t)† | Normalised TAO reflectance Landsat 7 band 5 (SWIR) from the latest valid pixels in year t | 0 - 255 |
| last_b70(t)† | Normalised TAO reflectance Landsat 7 band 7 (SWIR) from the latest valid pixels in year t | 0 - 255 |

* optional layer
† latest available cloud-free observation at pixel

We constructed an additional feature from Hansen et al. (2013) data called $recent\_loss(t)$. Deforestation tends to cluster around an emergent point (contagion) so we wanted to encode the proximity, in time and space, of recent loss. Therefore, we wanted to have feature that summarised the information of neighbouring pixels' deforestation state. We chose to represent recent loss as four, one-hot encoded layers, each layer representing loss within a specified period in the recent past.

All our models were build so that they can take two or more tensors with the same spatial dimensions, which we define below, and forecast if deforestation is observed in the following year at the locating corresponding to the spatially-central pixel of these tensors.

The first 3D tensor that any of our models receives, which we named **"Static"**, is tensor of shape $\mathbf{S} \in R^{2\times(2r+1)\times(2r+1)}$ where $(2r+1\times 2r+1)$ is its spatial dimension and $r$ is a predefined hyperparameter indicating the number of pixels the input tensor have in each spatial direction from the target central pixel (which has spatial coordinates $(r+1\times r+1)$ for an image of spatial size $(2r+1\times 2r+1)$ ). The two channels of this tensor are $treecover2000$ and $datamask$.

Our second set of tensors is a time series of 3D tensors $\mathbf{X}_{t-3}, \mathbf{X}_{t-2}, \mathbf{X}_t \in R^{5\times(2r+1)\times(2r+1)}$, where again each tensor has spatial dimensions $2\times(2r+1)\times(2r+1)$ but depth 5. The five channels of a tensor with time index $t$ are $recentloss(t)$, $last\_b30(t)$, $last\_b40(t)$, $last\_b50(t)$ and $last\_b70(t)$ as defined in Table S2.

Finally each tensor with time index $t$ comes with a label $Y_{t+1} \in \{0, 1\}$ which takes value 1 only if the target central pixel (at spatial location $r + 1 \times r + 1$) is marked as deforested exactly in year $t + 1$. To clarify this, here we note that if this pixel was labeled as deforested in any other year $t_j \neq t + 1$ ,$lossyear_{t_j} = 1$, or was never labeled as deforested in the study period 2001-2018 ,$lossyear_t = 0 \forall t_j in 1, 2, ..20$, then $Y_{t+1} = 0$

Due to the characteristics of Hansen et al. (2013) dataset, we know that if a pixel is labeled as deforested in year $t_j$ then the pixel never returns to the state of being forested. Additionally, if its the percentage of tree cover observed in 2000 was below 30%, than this location is not considered as forest. Only if a pixel with $treecover2000 < 30\%$ experience "gain" in the study period 2001-2012 we may assume it corresponds to a forested area from 2013 onward. Finally, if it has $datamask = 1$ then we know it is a permanent water body. Having stated this facts, we note that if our models aim to forecast the label of a pixel with index $j$ , $Y^j_{t+1} \equiv I\{lossyear_j = t + 1\}$, they would not be of any use if we know that this pixel j is not a forested area in year t. It will never be reverted to forest and therefore detecting deforestation at this location in year $t + 1$ doesn't make sense. Therefore, when predicting the labels of pixels ${Y^j}_{t+1}$ in year $t + 1$, we restricted these set of pixels to be:

$$\mathbf{J}_t =: \{j \in M : (lossyear_j > t \ \cup \ lossyear_j = 0)$$

$$\cap (datamask_j = 0) \cap (treecover_j > 30\% \ \cup \ gain_j = 1)\}$$

where $M$ is the index set of pixels lyng within Madre de Dios boundaries.

Since channel $treecover2000$ has range 0:100, and the Landsat bands 0:255, we rescaled each of them to be in the range 0:1.

For our last 3 models that utilize a time series of tensors we worked with the following dataset: $[\mathbf{S^j}, \mathbf{X}^j_{2014}, \mathbf{X}^j_{2015}, \mathbf{X}^j_{2016}]$ as set of input tensors and $Y^j_{2017}$ as the set of labels to be predicted where $j \in \mathbf{J}_{2016}$. We split the data into train,validation and test with ratio 6:2:2 to select the best model of each class. We evaluated their performance on $[\mathbf{S^j}, \mathbf{X}^j_{2015}, \mathbf{X}^j_{2016}, \mathbf{X}^j_{2017}]$ as the set of input tensors and $Y^j_{2018}$ as the set of labels to be predicted where $j \in \mathbf{J}_{2017}$.

Our Model 1, 2D CNN model, is able to analyze only mono-temporal tensors and from them to extract features forecasting the central pixel deforestation label in the following year. We used the union of the following data pairs of tensors and labels as dataset: $[\mathbf{S^j}, \mathbf{X}^j_{2014}]$ as an input tensors and $Y^j_{2015}$ as the set of labels to be predicted where $j \in \mathbf{J}_{2014}$. $[\mathbf{S^j}, \mathbf{X}^j_{2015}]$ as an input tensors and $Y^j_{2016}$ as the set of labels to be predicted where $j \in \mathbf{J}_{2015}$. $[\mathbf{S^j}, \mathbf{X}^j_{2016}]$ as an input tensors and $Y^j_{2017}$ as the set of labels to be predicted where $j \in \mathbf{J}_{2016}$. We evaluated its performance on : $[\mathbf{S^j}, \mathbf{X}^j_{2017}]$ as an input tensors and $Y^j_{2018}$ as the set of labels to be predicted where $j \in \mathbf{J}_{2017}$.

Here we note that to choose the best trained model form each model class, Model 2, Model 3, Model 4, we used as validation and text data that has labels in 2017, and therefore this models were biased towards the more recent year. Therefore, when choosing our best trained 2D CNN model, we use all data pairs to train it, but for early stopping validation data and model selection test data we used the pair $[\mathbf{S^j}, \mathbf{X}^j_{2016}]$ - $Y^j_{2017}$, $j \in \mathbf{J}_{2016}$. More details about our training strategy are provided in Chapter 6.

### S3 Model architectures

#### S3.1 Components

We recommend Goodfellow et al. (2016) for background on convolutional neural networks. Below is some key selected background material relevant to our model architectures.

##### S3.1.1 Convolutional Neural Network

Our data set was spatially organized into a grid with pixel-level observations. A leading type of Neural Network that are specially designed for analyzing grid-structured data is Convolutional Neural Networks (CNNs). CNNs have similarities with the general Artificial Neural Networks (ANNs) : they are still built up of filters that have learnable weights and biases and each of the filter takes some inputs, perform a dot product and activate it with a non-linear function. They differ in that they assume that the input data has a specific structure that determines their filters shape. Moreover, in each layer, the filters of CNNs are only linked with a local regions of the input tensor rather than with the full set of input's entries as are filters (neurons) in the fully connected layers of ANNs. Figure S1 illustrates how these two networks differ. Because in ANNs the outputs of the filters are scalar values, they are usually called neurons. In an ANN, a neuron $n$ in layer $l$ takes some input vector $\mathbf{x}$ (or the outputs of previous layer $l-1$), performs a dot product of all its values with its learnable weights $\mathbf{w}$ and add the bias term $b$. Then this neuron output is activated by a non-linear activation function $f$. Thus, a single neuron returns a scalar feature after processing all the input data. Any information about the structure of the input is lost. In CNNs each filter slides ("convolve") across one or more dimensions of the input tensor, performs a dot product between its entries and the input entries at these local regions and record its response at each location in an activation map. The number of dimensions across which the filter slides determines the dimensionality of the output map. With 1D-, 2D- or 3D-CNNs one usually refers to the number of dimensions across which the filters slide. In this project we used models that involve 2D and 3D convolutional filters. Below we present the key concepts of 2D and 3D CNNs that one needs to know to understand the architecture of our models.

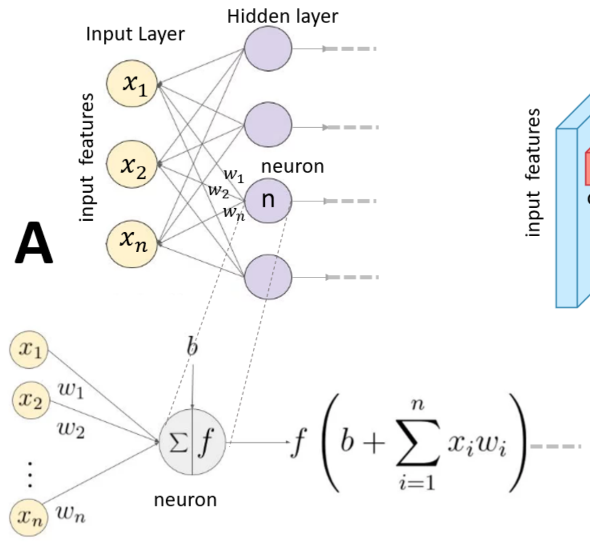

**Figure S1: A)** neuron(filter) in an Artificial Neural Network is connected to all neurons in the previous layer (input features). It preforms a dot product of its weights with the input feature vector and returns a scalar activation value. **B)** A filter in a 2D Convolutional Neural Network has 3D shape. It slides across the height and width of the input 3D tensor (features) and thus is connected to only one local region of the input at a time. It performs a dot product of its weights with the input entries at that location and map the activated value to a 2D map of activations (feature map). Adapted from: Fei-Fei Li (2017)

###### S3.1.2 2D Convolutions

In this subsection we present the key concepts of 2D

While a black and white image has a 2D rectangular shape, colour images are represented as a stack of three 2D grid maps each of which specify the intensity of Red, Green and Blue present at a particular location on the grid. The mixture of the three forms the colour. For hyper/multi-spectral images the number of channels is usually considerably larger. 2D CNNs for image processing constrain their filters to be made of set of spatially small 2D kernels, where the number of kernels always extends to the number of channels in the image. The stack of these kernels along the channel axis forms the 3 dimensional filter. The filter slides across the height and width of the input, performs a dot product at each location and its bias term and map its activated responses to a 2D feature map (activation map). Figure S2 illustrates this 2D covolutional operation. In the figure the image has size $5 \times 5 \times 3$ (height, width, depth). Additionally, a zero padding of size 1 is applied which makes its size $7 \times 7 \times 3$. Padding means to add extra pixels outside the image (zero padding is when these added pixels have value 0 in all of their channels and the size of the padding defines how many pixels are added in each direction). In Figure S2 two filters are present in the first hidden layer of the network. Each of them has weights with dimension $3 \times 3 \times 3$ and a bias term. The image and the filters are shown as sliced across the channel domain. Take the first filter for example. It performs a dot product at each location and maps it to a 2D feature map. The equation for 2D Convolution is as follows:

$$v_{l,j}^{x,y} = f\left(\sum_{m=0}^{M-1}\sum_{h=0}^{h=H_j-1}\sum_{w=0}^{w=W_j-1} k_{h,w}^{ljm} v_{(l-1),m}^{(x+h),(y+w)} + b_{lj}\right)$$

Where $l$ denotes the layer where the new output $v$ is. $j$ is the number of feature maps in this layer $l$ and $M$ is the number of feature maps (the depth of the input 3D tensor) in the previous $l-1$ layer. Applying j filters to an image results in output with j feature maps (3D output tensor of depth $j$). $x$ and $y$ are the spatial coordinates of $v$. $k_{h,w}^{ljm}$ is the $h, w, m$ value of the $j^{th}$ filter in layer $l$ and $b_{lj}$ is its bias term. $f$ is a non-linear activation function. In Figure S2 the highlighted output (=-2) is the $x, y = 1, 2$ entire of feature map $j = 1$ in layer $l = 2$. Here, the image is convolved at stride 2, where by stride 2 it is meant that the filter performs a dot products with local regions of the image that have their centers 2-pixels apart.

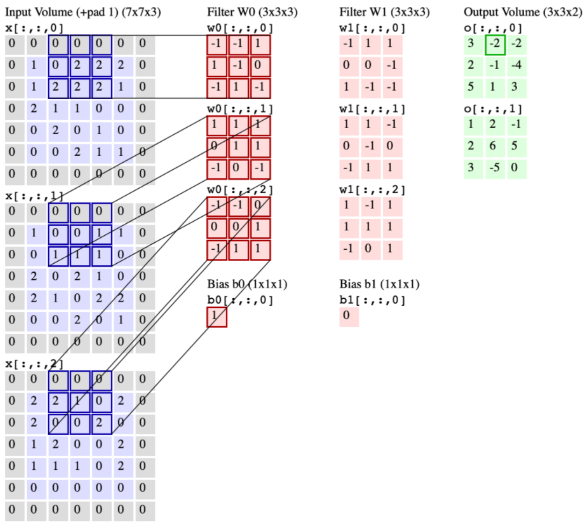

**Figure S2:** Convolution of 2 filters with an image. Sourced from: Fei-Fei Li (2017)

##### S3.1.3 3D Convolutions

In a 3D convolutional operation, filters slide across three of the dimensions of the input tensor and therefore they are made of a set of 3D kernels, where again this set size extends to the forth dimension of the input tensor. The stack of these 3D kernels makes the filter four dimensional tensor. As our data set has four dimensions: channels, time, height and width, if we slide the filters across the space and time domain, the network will be able to learn filters that get activated when they detect some type of spectral feature at particular time and location. 3D convolution is analogous to the 2D convolution with the difference being that the filter and the tensor has one more additional dimension. To visualize how 3D convolution works, consider Figure S3, where we present a four dimensional tensor as sliced across its forth dimension - the channel domain (the slices of the filter are denoted as 3D kernels). The filter slides along the three dimensions (height, width and time) of the input tensor, performs a dot product of its weights and the entries of the input tensor at these regions, add the bias and map the activated responses he gets at each spatio-temporal region to a 3D activation map. We present the equation for 3D convolution as Li et al. (2017) did. It is:

$$v_{l,j}^{x,y,z} = f\left(\sum_{m=0}^{M-1}\sum_{h=0}^{H_j-1}\sum_{w=0}^{W_j-1}\sum_{t=0}^{T_j-1} k_{h,w,t}^{ljm} v_{(l-1),m}^{(x+h),(y+w),(z+t)} + b_{lj}\right)$$

Where again $l$ denotes the layer where the new output $v$ is. $j$ is the number of 3D feature maps in this layer $l$ and $M$ is the number of feature maps in the previus layer (the forth dimension of the input 4D tensor). $x$,$y$ and $t$ are the spatio-temporal coordinates of $v$. $k_{h,w,t,m}^{ljm}$ is the $h, w, t, m$ weight of the $j^{th}$ filter ($\in R^{m\times H_j\times W_j\times T_j}$) and $b_{l,j}$ is its bias term. $f$ is a non-linear activation function. Applying j filters to a 4D tensor results in output with j 3D feature maps, which when stack together, form the new four dimensional tensor of activations.

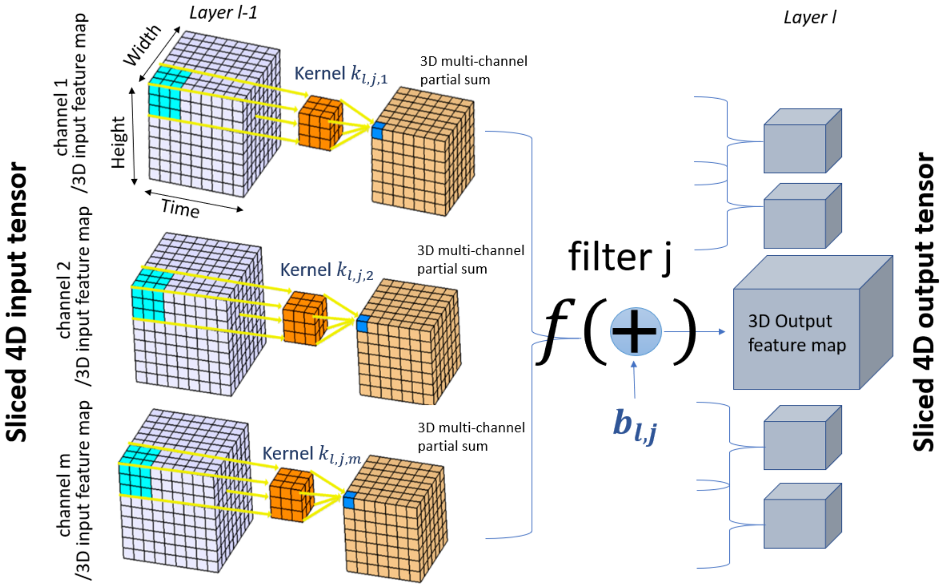

**Figure S3:** 3D Convolution of 4D tensor. Adapted from: Kaggle (2018)

##### S3.1.4 Deep Convolutional Neural Networks

Networks are "deep" when they have more than one hidden layer. All layers presented in the figures above ( Figure S1,S2 and S3) are the first hidden layer of the networks which take as an input the features of the input layer - the raw images. A layer is called hidden if it is not the input or the output layer. A network is called "Convolutional" if it has at least one convolutional filter in any of its layers. As we discussed above, in a 2D CNN each filter slides along 2 dimensions of the 3D input tensor and returns a 2D activation map. The stack of all filters' activation maps in a layer is the new 3D tensor of high level features that will be propagated to the next layer. For a 3D convolution, the 3D activation maps of the filters in a layer are stacked to form the new 4D tensor. By propagating the image in such a way through the layers, the network is able to extract high level features.

Figure S10 illustrates the architecture of our first proposed model, which is indeed a deep 2D Convolutional Neural Network. Our 2D CNN model takes as an input a mono-temporal multispectral image, propagates it through the network and returns a softmax output that indicates the probability of observing deforestation at the location where the center of the image is. Hereby we note once again, the task of this network is classification forecasting, rather than nowcasitng. In figure S10 one can see that each convolutial layer, activated with rectified linear unit (Hinton (2010) $ReLU(x) = max(x, 0)$), is followed by a layer called "2D Batch Normalization" layer. In the diagram of our model's architecture one can also see layers named as "Dropout" and "Spatial Pyramid Pooling Layer". In the rest of this section we explain what these layers do and discuss several issues one must consider when utilizing them.

##### S3.1.5 Batch Normalization

Batch Normalization was introduced by Ioffe, Szegedy (2015). It is a layer that normalizes each filter to have a zero mean and unit variance. Ioffe, Szegedy (2015) showed that employing such layers in Neural Networks can be beneficial in several ways: the networks train faster as it enables the gradient descent algorithm to take higher learning rates; the convergence of the loss function is significantly less sensitive to how the weights are initialized; it offers some level of regularization by adding small noise to the data and sometimes can even work as well as dropout which can decrease the need of dropout layers present in the network. A 2D Batch Normalization layer in a CNN as proposed by Ioffe, Szegedy (2015) normalizes the entries of each feature map before their activation. The normalization is done by taking the mean and variance estimated across all locations and batches. Consider a minibatch that has $m$ 3D tensors (height, width and depth) that are convolved with the $d$ filters of the current layer. The output of this layer is then a stack of $d$ 2D feature maps. During training each of $(l, k)^{th}$ element $(l \in 0, 2, ..H_j, k \in 0, 1..W_j)$ of the $j^{th}$ $(j = 1, 2, 3, , ..d)$ feature map that evolves from processing image under index $b$ $(b \in 0, 1, 2, 3...m)$ in the batch is transformed as follows:

$$\hat{y}_{b,j,l,k} = \hat{x}_{b,j,l,k} \times \gamma + \beta \qquad \hat{x}_{b,j,l,k} = \frac{x_{b,j,l,k} - \hat{E}[x_j]^{Moving}}{\sqrt{\hat{\sigma}_j^{Moving}} + \epsilon}$$

where $\gamma$ and $\beta$ are learnable scale and shift parameters and $\epsilon$ is a constant added for numerical stability. Also:

$$\hat{E}[x_j]^{Moving} = \sum_B \hat{E}[x_j] \qquad \hat{\sigma}_j^{Moving} = \frac{m \times H_i \times M_i}{m \times H_i \times M_i - 1} \sum_B \hat{\sigma}[x_j]$$

Are the moving averages of the empirical mean($\hat{E}[x_j]$) and variance ($\hat{\sigma}^2[x_j]$) across batches ($B$ is the batches index set). During inference they are kept as the constants.

$\hat{E}[x_j]$ and $\hat{\sigma}^2[x_j]$ are obtained as follows:

$$\hat{E}[x_j] = \sum_{i=0}^{m}\sum_{h=0}^{H_j}\sum_{w=0}^{W_j} x_{i,j,h,w} \quad \hat{\sigma}^2[x_j] = \frac{1}{m_{jj}}\sum_{i=0}^{m}\sum_{h=0}^{H_j}\sum_{w=0}^{W_j}(x_{i,j,h,w} - \hat{E}[x_j])^2$$

Where $H_j, W_j$ are the spatial size of the $j^{th}$ feature map and $x_{i,j,h,w}$ is the $(h,w)^{th}$ entire of the $j^{th}$ feature map of the $i^{th}$ 3D tensor in the batch. The equations for 3D Batch Normalization is analogous with the difference being in that feature maps are 3D tensors and hence:

$$\hat{E}[x_j] = \sum_{i=0}^{m}\sum_{t=0}^{T_j}\sum_{h=0}^{H_j}\sum_{w=0}^{W_j} x_{i,j,t,h,w} \quad \hat{\sigma}^2[x_j] = \frac{1}{m \times T_j \times H_j \times M_j}\sum_{i=0}^{m}\sum_{t=0}^{T_j}\sum_{h=0}^{H_j}\sum_{w=0}^{W_j}(x_{i,j,t,h,w} - \hat{E}[x_j])^2$$

In their original paper Ioffe, Szegedy (2015) employed Batch Normalization by first normalizing the entries of the feature map $\hat{x}_{b,j,l,k} \to BN[x_{b,j,l,k}] = \hat{y}_{b,j,l,k}$ and then activating them by a non-linear activation function $f(\hat{y}_{b,j,l,k})$ , e.g. ReLU. However, the case what should be the right order of applying $f$ and $BN()$ to the input features is a topic of debate. Although up to our knowledge there isn't a scientific body of work that address this problem, as many other experts in the field, we empirically showed that when ReLU is applied before a Batch Normalization layer : $y = BN[ReLU(x)]$, our networks perform better. The results of these experiments are shown in the Appendix.

##### S3.1.6 Dropout

Dropout was proposed by Srivastava et al. (2014) as a technique for regularizing neural networks by adding noise to the entries of the hidden layer. More precisely, during training it works by multiplying the hidden activations with a Bernoulli random variable which takes value 0 with probability p or 1 - with 1-p respectively. As networks get deeper, the number of weights grows exponentially. This cause networks to overfit if no regularization measurements are employed. Employing dropout approximates an inexpensive way of training and inference of exponentially many networks. The way this is done is by randomly switching off different neuron units during each training forward pass. Without some of its neurons, the network represents a different function, or sub-network. When trained with dropout, the network cannot depend on any given neuron as it might be suddenly dropped out. This prevents it from learning features that depends on each other and also from returning an output that depends on one particular feature. During inference, we want the model to use all of its learned weights and not to drop out. When deployed, we multiply the scores of each neuron by the probability of it not being dropped $1 - p$. To understand how it works, let us consider a feature vector of activations $\mathbf{x} = (x_1, x_2, ..x_d)$. If we apply dropout on this layer, the vector becomes $\mathbf{x} = (a_1x_1, a_2x_2, ...a_dx_d)$ where $a_k$ are independent Bernoulli random variables. When testing, the vector becomes $\mathbf{x} = ((1-p) \times x_1, (1-p)x_2, ...(1-p) \times x_d)$ (See figure S4). Alternatively, one can scale up the activations by multiplying them with $\frac{1}{1-p}$ during training and not modify them at inference (this is how pytorch implements it). In our models we have employed dropout in the second to last fully connected layer. The reason for that is because each of other layer of our networks is followed by a Batch Normalization layer. In their research Li et al. (2018) theoretically showed that although both Batch Normalization layer and Dropout layer are very powerful regualization tools, their joint utilization in a neural network can lead to worse performance. The reason for this is because employing a Dropout layer has the side effect of shifting the variance of the neurons when the model is switched form training to inference state. On the other side, at inference the Batch Normalization layer maintains its statistical variance that has been learned during training. This variance dismatch, which Li et al. (2018) defined as "variance shift", makes the model unstable if a Dropout layer is applied before a Batch Normalization layer. Li et al. (2018) also confirmed their findings empirically by performing experiments on widely used networks architectures. The theory about these"variance shift" can be find in the Appendix. Li et al. (2018) then suggested two methods that can prevent this variance disharmony when both regularization techniques are used: the simpler one is Dropout layer to be applied only after the last Batch Normalization layer, and the other is to use modified formula for the Dropout scaling factor 1-p. In our model we employed their first suggestion and only used dropout after the last Batch Normalization Layer. We confirmed their suggestion is valid by performing experiments on applying Dropout layer in several other layers, which were then followed by a Batch Normalization layer. Results of these experiments can be find in the Appendix.

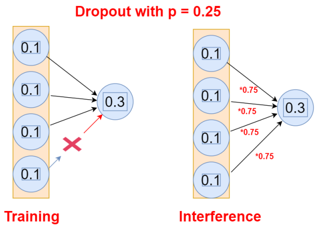

**Figure S4:** Applying Dropout in a fully connected layer with probability of dropping a neuron p = 0.25. During inference each feature is multiplied by 1-p = 0.75.

##### S3.1.7 Spatial Pyramid Pooling layer

The final piece of architecture present in the first two models that has not yet been discussed is the Spatial Pyramid Pooling layer. To begin with, general pooling layers are another type of layers commonly used in 2D CNNs architectures that are inserted between convolutional layers. However, their filters do not have learnable weights. Their only function is to progressively decrease the input spatial size which consequently decrease the number of network parameters. The effect of employing pooling layers is reduced number of computational operations and decreased overfitting. Pooling layers vary depending on the way they reduce the impute size: max-pooling layers, average-pooling layers, etc. The pooling filters slide along the image and independently downsample each depth slice of the input. Figure S5 demonstrate how a Maxpool filter of spatial size $2 \times 2$ slides across the input height and width at stride 2 and return the downsampled 3D tensor with decreased spatial size by factor of 2.

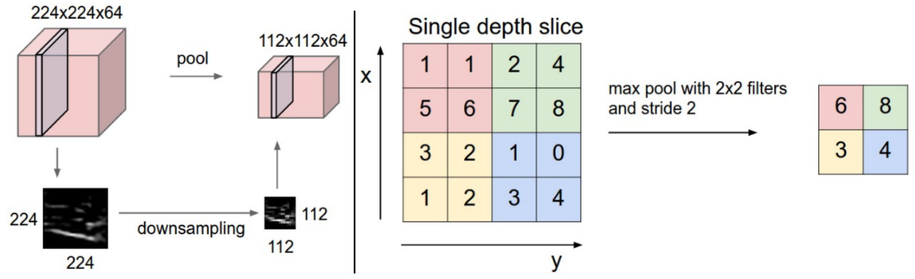

**Figure S5:** Independently downsamling each slice of an input volume with a Maxpooling filter of spatial size$2 \times 2$ at stride 2. **Left:** Input volume of size $224x224x64$ result in a volume of $112x112x64$ (Note the depth of the volume is preserved). **Right:** Downsampling of a single slice with the corresponding Maxpool filter's kernel. The $2 \times 2$ kernel slides across the input slice and returns the maximum pixel value it receives at each location. Source: Fei-Fei Li (2017)

Standard 2D CNN architectures for image classification usually consist of several to many convolutional layers which are then followed by fully connected layers. The last high-level 3D feature map generated by the final convolutional layer is then flattened to a 1d vector and passed to the first fully connected layer. The size of this 1d feature vector is determined by the size of the final 3D feature map which also defines the number of filters the network has in the first fully connected layer. While the number of parameters in the convolutional layers depends only on the filters' sizes, the output of these layers depend on the filters and image sizes. Therefore in order to be able to design an architecture that has convolutional followed by fully connected layers, the size of the image must be predefined. He et al. (2014) proposed another pooling strategy, "Spatial Pyramid Pooling", that enables networks to be trained on images with various spatial sizes. They proposed 2D CNN with a spatial pyramid pooling layer between the last convolutional and the first fully connected layer, called "SPP-net", which has fixed number of parameters and is able to analyze images regardless of their spatial size. They demonstrated that employing the Spatial Pyramid Pooling strategy improve on many widely used CNNs architectures for image classification and object detection tasks that fit the input image to the required size by cropping or padding it. We decided to implement their strategy as we wanted to assess our model's performance when the input satellite images capture smaller or larger land regions without the need of prepossessing the images or modifying the models. This also allowed us to explore what is the optimal image size for a model with fixed number of parameters. Figure S6 illustrate how the Spatial Pyramid Pooling layer takes a 3D tensor with arbitrary spatial size and returns a fixed-size feature vector.

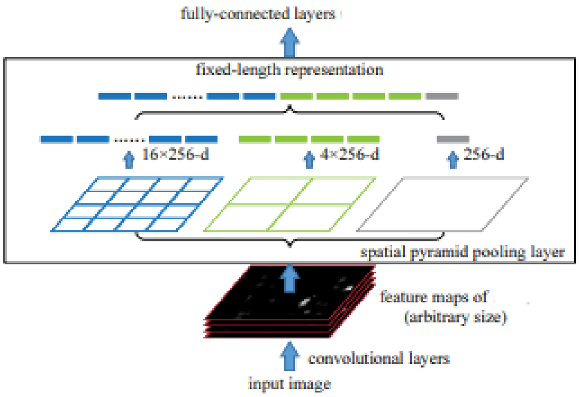

**Figure S6:** Input 3D tensor of arbitrary spatial size and depth 256 is forwarded to a spatial pyramid pooling layer that has 3 filters. The three filters slide across the spatial domain of the input and return 3D tensors of size $16 \times 16 \times 256$ , $4 \times 4 \times 256$ and $1 \times 1 \times 256$ respectively. There tensors are then flattened and connected together to form a feature vector of fixed size = $256 \times (16 \times 16 + 4 \times 4 + 1 \times 1)$ that is then the input of the following fully connected layer.Source: He et al. (2014)

A Spatial Pyramid Pooling layer has predefined fixed number of pooling filters. One can chose the function with which these filters downsample: max, average, etc. However, the spatial size of these filters and the stride at which they slide across the spatial domain of the input tensor is dynamic, it adapts according to the spatial size of the input. This filters are then able to return 3D tensors of predefined spatial size and same depth as the input tensor. Having fixed number of 3D output tensors with fixed spatial size and depth, when flattened and contacted together, they form a fixed-size feature vector. In Figure S6 the spatial pyramid pooling layer has predefined number of filters 3 with predefined spatial sizes $16 \times 16$, $4 \times 4$ and $1 \times 1$ respectively. The input 3D tensor has arbitrary spatial size and depth of 256. After flattening and connecting the output of each pooling filter, the extracted 1d feature vector has size $256 \times (16 \times 16 + 4 \times 4 + 1 \times 1)$. The equations that govern the dynamics of the poling filters' spatial sizes and stride is as follows:

$s_h = floor(H_x/H_k)$ $\qquad$ $s_w = floor(W_x/W_k)$

$k_h = floor(H_x/H_k) + H_x mod(H_k)$ $\qquad$ $k_w = floor(W_x/W_k) + W_x mod(W_k)$

Where $s_h, s_w$ are the strides at which the pooling filter must slide across the height $H_x$ and width $W_x$ of the input 3D tensor. This filter must also have height and width $k_h$ and $k_w$ respectively. The output after sliding it across the input tensor has the desired height and width $H_k, W_k$ and same depth $d$ as the input tensor depth.

###### S3.1.8 Recurrent Neural Networks

Rather than learning features that can explain the current state of a pixel, our model differs in that it tries to learn features that can identify if the target area will become deforested the following year. Our hypothesis is that those features can be extracted from past and current satellite images and moreover, that the temporal structure of these observations contains information that can be beneficial for the accuracy of our prediction - e.g if we assume that a deforestation event is not a sudden event but rather a continuous process, the model may be able to learn the target area's deforestation tendency.

There are two ways a model can learn these features - one is called feed forward and the other - recurrent. So far we presented our CNN model and 3D-CNN model as feed forward learning models - the signals flow one direction only: from the input layer of image features to the final layer producing the output (class label $\hat{Y}_{t+1}$). In such models the output of any layer does not affect the current or any of the previous layers. In our second model, Model 2: 3D CNN, we extract features $\mathbf{s}_{t+1}$ given a 4D input tensor that is formed by concating sequence of inputs images $\mathbf{x}_{t-k}, \mathbf{x}_{t-(k-1)}, .., \mathbf{x}_t$ with a fixed window size $k$ along the time axis and propagate it forward through the network until the final layer that has softmax activation indicating our confidence of the central pixel label.

In a recurrent learning process, the signals may travel both forward and backward direction by introducing loops in the network. Figure S7 outline the main difference between feed-forward and recurrent learning networks. The forward propagation of a simple "vanilla" Recurrent Neural Network with only one hidden layer is governed by the following equation:

$$\mathbf{s}_t = \phi(\mathbf{W}_s \times \mathbf{s}_{t-1} + \mathbf{U}_x \times \mathbf{x}_t + \mathbf{b}) = \phi([\mathbf{W}|\mathbf{U}] \times [\mathbf{s}_{t-1}|\mathbf{x}_t]^T + \mathbf{b})$$

Where:
$\times$ is matrix multiplication operation,
$\mathbf{s}_i \in R^h$, $\mathbf{x}_i \in R^m$, $\mathbf{W} \in R^{h \times h}$, $\mathbf{U} \in R^{h \times m}$, $\mathbf{b} \in R^h$,
so that $[\mathbf{W}|\mathbf{U}] \in R^{h\times(h+m)}$ and $\mathbf{b} \in R^h$ are the weight matrix and the bias vector of the hidden layer and $[\mathbf{s}_{i-1}|\mathbf{x}_i]^T \in R^{h+m}$ is its input. $\phi$ is a non-linear squashing function, usually hyperbolic tangent. We set $\mathbf{s}_0 = \mathbf{0}_h$.
We see that in thus defined forward propagation, the output of a hidden layer $\mathbf{s}_{k-1}$ is looped back and contacted with the new input image $\mathbf{x}_k$. This new input ($[\mathbf{s}_{t-k}|\mathbf{x}_k]^T$) is then passed again to the same hidden layer and a new "updated" $\mathbf{s}_k$ output is returned, allowing the network to process the images sequentially. The iteration continues until all input images are learned. At each step, the output $\mathbf{s}_k$ has "learned" all the input images the network have seen so far: $\{\mathbf{x}_{k-1}, \mathbf{x}_{k-2}.. \})$. Due to this internal memory, RNN's are able to remember important information about the new inputs they received, which makes them powerful in predicting what's coming next.

This ability to "memorize" have made them very popular in tasks that deals with sequential data type. Depending on the objective, several types of architectures have been developed. One possible architecture is "many-to-many" where at each iteration the model use the most recent $\mathbf{s}_k$ to predict the future output (in our case this can be class label $\hat{Y}_{k+1} = \theta(\mathbf{s}_k)$). In this case the loss is evaluated as sum of all individual losses: $L_{tot} = \sum_k L_k = BCEloss(\hat{Y}_{k+1}, Y_{k+1})$. Other tasks may require a single prediction of a future event, given past time-series data, for which one usually use the architecture "many-to-one". In such a model, we may only use the final "most knowledgeable" feature vector $\mathbf{s}_t$ to make a prediction and may only report the final loss $L = BCEloss(\hat{Y}_{t+1}, Y_{t+1})$. This two differences are reflected in the red square of Fig S7 **B)**. Due to their empirically proven power, many variations of RNNs have been developed.

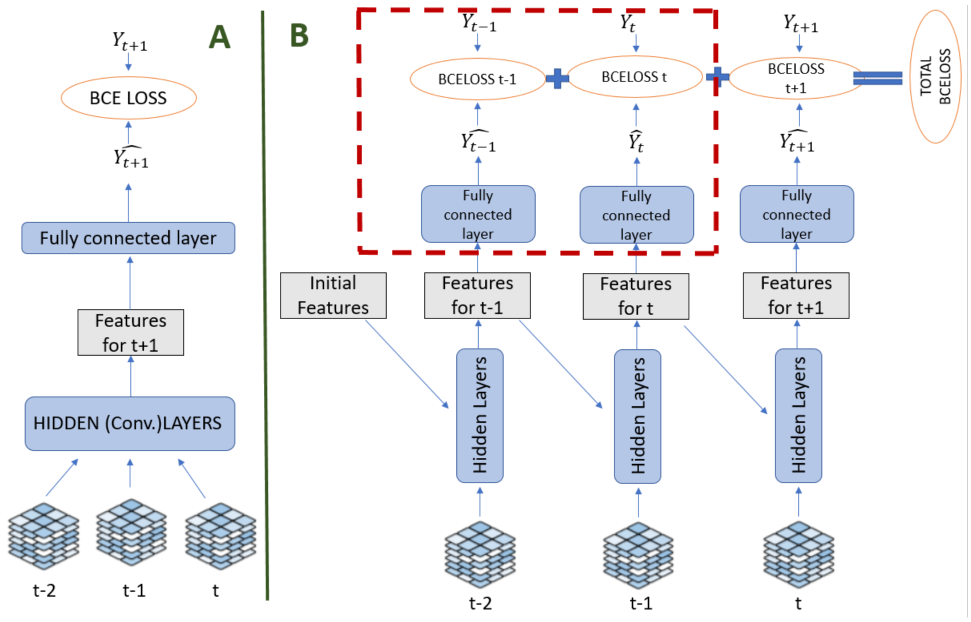

**Figure S7: A)** CNN model with feed forward architecture: The signals from the images travel one direction only,from the input layer to the output layer; **B)** Vanilla Recurrent Neural Network has recurrent learning: the output of the hidden layer is looped back, cocated with the new image and forwarded to the same layer again. When the output of the layer is also forwarded to the next layer at each time, the model has "many-to-many" architecture. When the hidden layer forwards the output only at the last time itteration, the architecture is "many-to-one".

##### S3.1.9 Long Short Term Memory Cell

Nevertheless, when dealing with long sequential data, during learning, back- propagation, a "Vanilla" RNN may suffer from exploding or vanishing gradients. While for the first case, exploding gradients, truncating the gradients can solve the undesirable effect, the vanishing gradients problem is much harder to overcome. Fortunately,Hochreiter, Schmidhuber (1997) proposed the concept of Long-Short Term Memory RNN structure which was able to solve the vanishing gradients problem. Since than, several variations of the LSTM RNNs were proposed. Below we briefly outline the concept of the LSTM cell as defined in the Deep learing book by Goodfellow et al. (2016). The governing equations in an LSTM cell are as follows:

$$\mathbf{f}_t = \sigma(\mathbf{W}_f\mathbf{h}_{t-1} + \mathbf{U}_x\mathbf{x}_t + \mathbf{b}_f) = \sigma([\mathbf{W}_f|\mathbf{U}_f] \times [\mathbf{h}_{t-1}|\mathbf{x}_t]^T + \mathbf{b}_f)$$
$$\mathbf{i}_t = \sigma(\mathbf{W}_i\mathbf{h}_{t-1} + \mathbf{U}_i\mathbf{x}_t + \mathbf{b}_i) = \sigma([\mathbf{W}_i|\mathbf{U}_i] \times [\mathbf{h}_{t-1}|\mathbf{x}_t]^T + \mathbf{b}_i)$$
$$\mathbf{o}_t = \sigma(\mathbf{W}_o\mathbf{h}_{t-1} + \mathbf{U}_o\mathbf{x}_t + \mathbf{b}_o) = \sigma([\mathbf{W}_o|\mathbf{U}_o] \times [\mathbf{h}_{t-1}|\mathbf{x}_t]^T + \mathbf{b}_o)$$
$$\mathbf{g}_t = \tanh(\mathbf{W}_g\mathbf{h}_{t-1} + \mathbf{U}_g\mathbf{x}_t + \mathbf{b}_g) = \tanh([\mathbf{W}_g|\mathbf{U}_g] \times [\mathbf{h}_{t-1}|\mathbf{x}_t]^T + \mathbf{b}_g)$$

$\mathbf{c}_t = \mathbf{f}_t \odot \mathbf{c}_{t-1} + \mathbf{i}_t \odot \mathbf{g}_t$
$\mathbf{h}_t = \mathbf{o}_t \odot \tanh(\mathbf{c}_t)$
$\times$ : matrix multiplication operator
$\odot$ : the Hadamard (elemnt-wise) product
$\mathbf{x}_i \in R^d$: input vector to the LSTM unit
$\mathbf{h}_i \in R^h$: output vector of the LSTM unit (also known as hidden state)
$\mathbf{g}_i \in R^h$: input activation vector
$\mathbf{c}_i \in R^h$: LSTM internal state vector
$\mathbf{f}_i \in R^h$: forget gate's activation vector
$\mathbf{i}_i \in R^h$: input gate's activation vector
$\mathbf{o}_i \in R^h$: output gate's activation vector
$\mathbf{W} \in R^{h\times h}$, $\mathbf{U} \in R^{h\times d}$ and $\mathbf{b} \in R^h$: weight matrices and bias vector parameters which need to be learned during training.

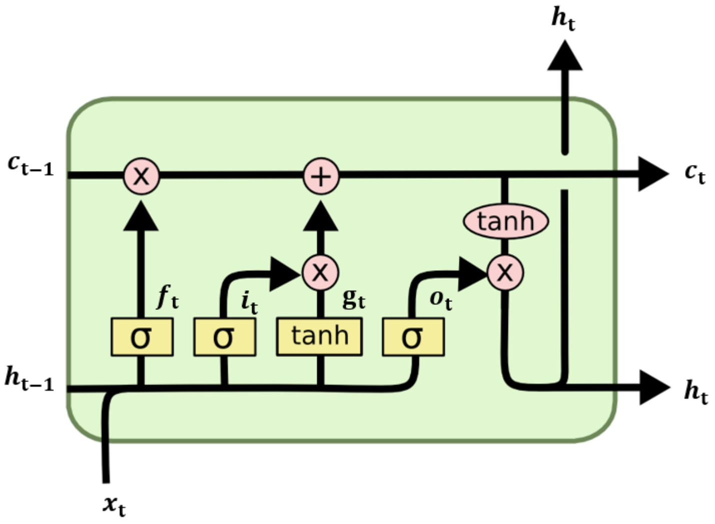

**Figure S8:** Information flow in a LSTM cell. Sourced from: Hoffman (2018)

Figure S8 illustrate this LSTM cell architecture. A new feature vector $\mathbf{g}_t$ is learned as in the "vanilla" RNN case - a regular neuron units (with weights matrix and bias vector $\mathbf{W}_\mathbf{g}|\mathbf{U}_\mathbf{g}|\mathbf{b}_\mathbf{g}$) are fed with the connected new input vector $\mathbf{x}_t$ and the previous output vector $\mathbf{h}_{t-1}$. In the case above, we have used hyperbolic tangent function to squash the neurons' outputs but any squashing nonlinear function can be used. In a LSTM architecture, however, this combination of present input and past output is also fed to three other gate vectors, which will decide how the new learned feature vector $g_t$ will be handled: they block, pass or output the new information based on its importance, which they have filtered with their own set of weights. How much of $\mathbf{g}_t$ will be accumulated in the internal state cell $\mathbf{c}_t$ is controlled by the input gate vector $\mathbf{i}_t$. A forget gate vector $\mathbf{f}_t$ decides how much of the current cell state information $\mathbf{c}_{t-1}$ will be blocked. Finally, the new output, the updated memory $\mathbf{c}_t$, is squashed with a hyperbolic tangent function but before being returned, it is regulated by the output gate vector $\mathbf{o}_i$. All the gate unit vectors have a sigmoid nonlinear activation, or also called squashing, function. Some variations of the LSTM architecture use the state cell as an extra input to the gate units. By temporally stacking several LSTMs layers above one another, a deeper network can be achieved.

##### S3.1.10 Convolutional Long Short Term Memory Cell

We now extend this idea of LSTM cell with the recently developed Convolutional LSTM structure. The modification of this LSTM RNN architecture as Convectional was first proposed by SHI et al. (2015). In the same year Ballas et al. (2015) proposed Convolutional RNN with Gated Recurrent unit (unit similar to the LSTM unit). The goal of SHI et al. (2015) work was to develop an algorithm that gives precise prediction of rainfall intensity in a local region over a short period of time, a problem which they defined as "spatiotemporal sequence forecasting". They used a sequence of past radar maps as an input and a sequence of a fixed number future radar maps as an output, task very similar to ours. Furthermore, they evaluated the performance of their model, which they named ConvLSTM, on the Moving-MNIST dataset and showed that ConvLSTM is able to achieve state-of-art accuracy on any task that deals with spatio-temporal sequence forecasting. Their design differs from the general LSTM cell in that all the inputs $\mathbf{x_1}, \mathbf{x_2}, ...\mathbf{x_t}$, hidden states $\mathbf{h_1}, \mathbf{h_2}, ...\mathbf{h_t}$, internal state cells $\mathbf{c_1}, \mathbf{c_2}, ...\mathbf{c_t}$ and gates of ConvLSTM are 3D tensors whose last two dimensions define their spatial dimensions. One can imagine them as vectors standing on a spatial grid. For each cell in that grid the ConvLSTM determines its future state and output by analyzing the input features and the past states of its local neighbours. This is achieved by substituting the matrix multiplication operations with convolutions. Below we provide the key equations of the ConvLSTM cell that we emplyed in our model. We adopted the ConvLSTM equations from Rußwurm, Körner (2018) and make note that they differ form those proposed by in that SHI et al. (2015) LSTM cell structure uses the state cell as an extra input to the gate units.

$\mathbf{f}_t = \sigma(\mathbf{W}_f * [\mathbf{h}_{t-1}|\mathbf{x}_t] + \mathbf{b}_f)$
$\mathbf{i}_t = \sigma(\mathbf{W}_i * [\mathbf{h}_{t-1}\mathbf{x}_t] + \mathbf{b}_i)$
$\mathbf{o}_t = \sigma(\mathbf{W}_o * [\mathbf{h}_{t-1}|\mathbf{x}_t] + \mathbf{b}_o)$
$\mathbf{g}_t = \tanh(\mathbf{W}_f * [\mathbf{h}_{t-1}|\mathbf{x}_t] + \mathbf{b}_g)$
$\mathbf{c}_t = \mathbf{f}_t \odot \mathbf{c}_{t-1} + \mathbf{i}_t \odot \mathbf{g}_t$
$\mathbf{h}_t = \mathbf{o}_t \odot \tanh(\mathbf{c}_t)$
$*$ denotes the convolution operator
$\odot$ denotes the Hadamard (elemnt-wise) product
$d, w, h$: the depth, width and height of the input tensor
$r$: number of channels of the hidden state, internal state cell and all the gates tensors
$k_1 \times k_2$: kernel size of the feature maps
$\mathbf{x}_i \in R^{d\times w\times h}$: input tensor to the LSTM unit
$\mathbf{h}_i \in R^{r\times w\times h}$: output tensor of the LSTM unit (hidden state)
$[\mathbf{h}_{i-1}|\mathbf{x}_i] \in R^{(r+d)\times w\times h}$: connected input tensor
$\mathbf{g}_i \in R^{r\times w\times h}$: input activation tensor
$\mathbf{c}_i \in R^{r\times w\times h}$: LSTM internal state tensor
$\mathbf{f}_i \in R^{r\times w\times h}$: forget gate's activation tensor
$\mathbf{i}_i \in R^{r\times w\times h}$: input gate's activation tensor

$\mathbf{o}_i \in R^{r\times w\times h}$: output gate's activation tensor
$\mathbf{W} \in R^{r\times(r+d)\times k_1\times k_2}$ are the weights of the $r$ stacked convolutional featues maps (each $\in R^{(r+d)\times k_1\times k_2}$) and $\mathbf{b} \in R^r$ is a vector with the bayes each of the $r$ features maps has. The initial state cell $\mathbf{c_0}$ and $\mathbf{h_0}$ are initialized as zero value tensors. To preserve the spatial size of the convolutional outputs we used zero padding of size $[\frac{k_1}{2}] \times [\frac{k_2}{2}]$ where we used kernels with odd sizes $k_1, k_2$.

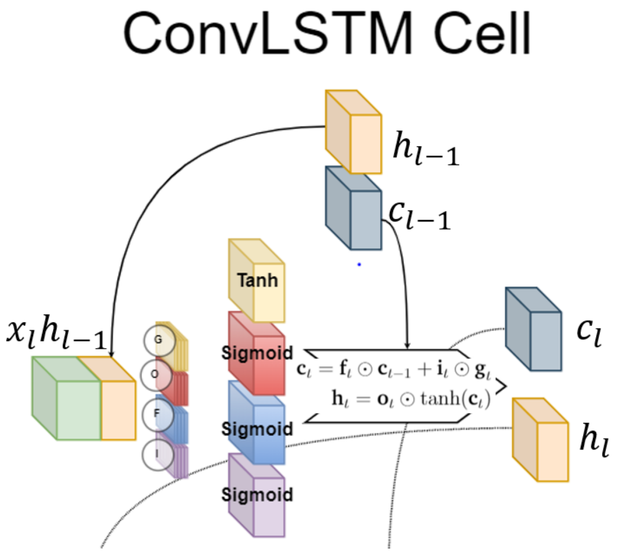

**Figure S9:** Convolutional LSTM Cell

#### S3.2 Model Architecture 1: 2D Convolution Neural Network

Our first proposed model, which we consider as our base model due to its simple architecture, is a 2D Convolution Neural Network with four 2D convolutional layers (2D Conv), one Spatial Pyramid Pooling layer (SPP) and 2 fully connected layers (FC) with a Dropout layer(DO) in between them. The final fully connected layer takes as input feature vector of size 100 and returns a scalar value squashed by a sigmoid non-linearity function, $\sigma$. The output is therefore in range [0,1] and indicates the model confidence of observing deforestation at the location corresponding to the spatially central pixel (with spatial cordinates $(r+1, r+1)$) of the input tensor in the following year. The filters of each layer are activated with ReLU and this activations are then normalized by a Batch Normalization layer(BN).

For a datapoint $j$, the input of this model is a 3D tensor, $\mathbf{SX}_t^j$, that dimension $R^{7\times(2r+1)\times(2r+1)}$. This input results from stacking its static 3D tensor, $\mathbf{S^j} \in R^{2\times(2r+1)\times(2r+1)}$, and one 3D tensor, $\mathbf{X_t^j} \in R^{2\times(2r+1)\times(2r+1)}$, of $j^s$ time series of 3D tensors $\{\mathbf{X_k^j}\}_{k=t-2}^{t}$. They are stacked along the channel axis so that the input of the network becomes $\mathbf{SX}_t^j$. The predicted label is $\hat{Y}_{t+1}^j$ where $j \in \mathbf{J}_t$ as defined in Chapter 4. Due to the SPP layer our model is capable of analyzing 3D tensor of any spatial size, which is regulated by $r$.

We have constructed this model such that we can vary the spatial size, the stride and the padding of the filters of any of the 2D convolutional layers (2DConv). The number of filters per convolutional layer is also set as free model parameter. The Spatial Pyramid Pooling layer (SPP) parameters are also allowed to vary, the number of pooling filters ,$n$, and the spatial size of this filters ,$\mathbf{k} \in R^{2,n}$ (see Chapter 2.2.3 for more details). By varying them one changes the size of the first fully connected layer (FC), as its size depend on the Spatial Pyramid Pooling layer output. Finally, the last free model parameter is the Dropout rate, p, with which the Dropout layer(DO) switches-off some of the connections between the two fully connected layers while training. As suggested by Li et al. (2018) we only employed a dropout layer after the last Batch Normalization Layer (BN). In this model architecture each convolutional layer is activated by ReLU, then these activations are normalized with a 2D Batch Normalization layer. The normalized features are then propagated to the next convolutional layer ($2DConv \rightarrow ReLU \rightarrow 2DBN \rightarrow 2DConv$). The normalized activations of the last 2D convolutional layer are then downsampled by the SPP layer and passed to the first FC layer. Its output of size 100 is activated again with ReLU and normalized with 1D BN layer. Only then dropping is implemented. We summarize the propagation of this model as follows: $\mathbf{SX}_t^j \in R^{7\times(2r+1)\times(2r+1)}$ $\rightarrow 4\times$(2DConv$\rightarrow$ReLU$\rightarrow$2DBN) $\rightarrow$ SPP($n, \mathbf{k}$) $\rightarrow$ FC(spp,100) $\rightarrow$ ReLU $\rightarrow$ 1DBN $\rightarrow$ DO(p) $\rightarrow$ FC(100,1) $\rightarrow \sigma \rightarrow \hat{Y}_{t+1}^j$

Figure S10 illustrates these architecture.

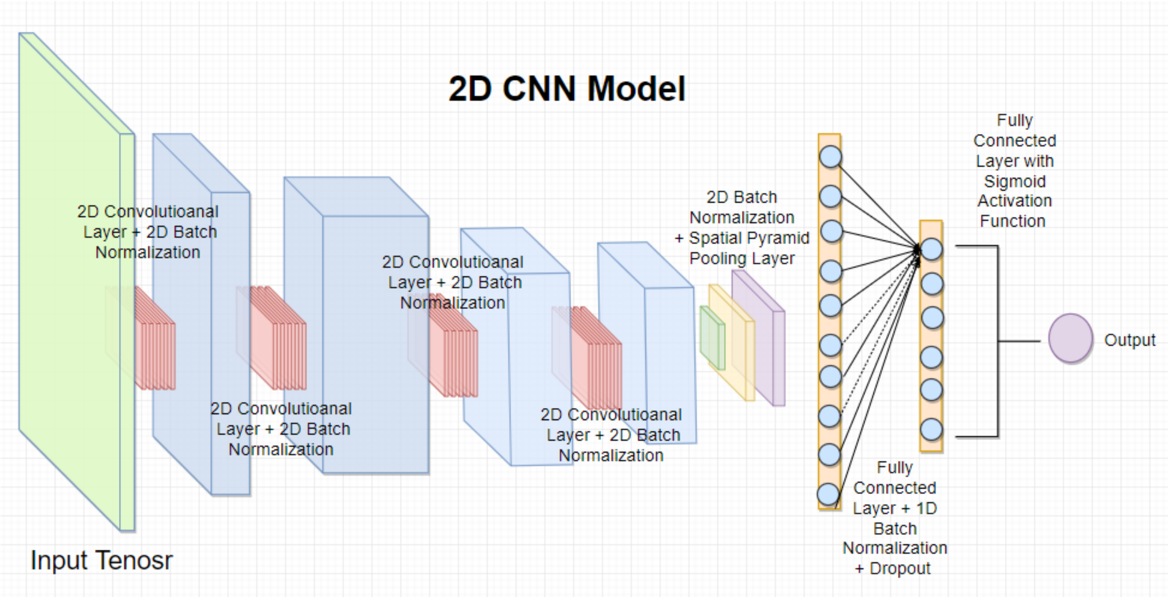

**Figure S10: Model 1:** 2D CNN. Architecture summary:
Input tensor $\mathbf{SX}_t^j \in R^{7\times(2r+1)\times(2r+1)}$ $\rightarrow 4\times$(2DConv$\rightarrow$ReLU$\rightarrow$2DBN) $\rightarrow$ SPP($n, \mathbf{k}$) $\rightarrow$ FC(spp,100) $\rightarrow$ ReLU $\rightarrow$ 1DBN $\rightarrow$ DO(p) $\rightarrow$ FC(100,1) $\rightarrow \sigma \rightarrow \hat{Y}_{t+1}^j$

#### S3.3 Model Architecture 2: 3D Convolution Neural Network

Our first proposed model is able to take advantage of the spatial and spectral domain of the input due to its 2D convolutional layers. However, it is not able to utilize the time domain of a datapoint $j$. It analyzes a 3D tensor $\mathbf{SX_t}^j$ that arses from stacking the Static features of $j$, $\mathbf{S}^j$, and only one element of $j^{th}$ time series $\mathbf{X}_t^j$.

Our second model, 3D CNN, takes on the other side takes full advantage of the time domain and thus is able to analyze $j^{th}$ spectral,spatial and time domain simultaneously. This is achieved via 3D convolutional layers present in its architecture, that consists of two branches which meet together to return the final prediction.

More precisely, for the $j^{th}$ data point the model takes as an input the $j^s$ static tensor $\mathbf{S}^j \in R^{2\times(2r+1)\times(2r+1)}$ and its full time series of non static tensors $\{\mathbf{X}_{t-2}^j, \mathbf{X}_{t-1}^j, \mathbf{X}_t^j\}$, each $\in R^{5\times(2r+1)\times(2r+1)}$. It then propagate these two parts of the input to two different branches. The static tensor $\mathbf{S}^j$ is passed to a 2D Convolutional Branch and the time series of tensors $\{\mathbf{X}_k^j\}_{k=t-3}^t$ to a 3D Convolutional Branch. Each branch extract high-level features which are then propagated together in the rest of the network. Finally, the model returns an output $\hat{Y}_{t+1}^j$ indicating the confidence of the model to observe deforestation at the target location in the following year t+1 where $j \in \mathbf{J}_t$ (see Chapter 4 for more details about ths notation).

Here we give more detailed explanation what each branch does. $\{\mathbf{X}_k^j\}_{k=t-3}^t$ is passed to a 3D Convolutional Branch in the form of 4D tensor obtained by stacking the sequence by the time domain. Thus the input tensor $\mathbf{X}^j \in R^{5\times3\times(2r+1)\times(2r+1)}$ has its first domain defined by the channels, the second by the time and the last two, by the space. The 3D Convolutional Branch "convolve" with $\mathbf{X}^j$ across its last three dimensions, time, height and width in two sequential 3D convolutional layers. Due to our limited time domain, of size 3, the 4D filters have shape $\in R^{chanels,2,k_h,k_w}$, where the first dimension extends to the number of the input channels and the last three define the shape of its 3D kernels. While setting different values to their spatial sizes is possible, the size of its time domain could only be 2. The model slides its filters' 3D kernels along thee time domain at stride 1 and no padding is applied on the input 4D tensor. Therefore, after the two 3D convlutional layers the output of the 3D Convolutional Branch was a 3D tensor of high-level features with no time domain, $\mathbf{Z}_x^j$. We summarize this propagation as follows: $\mathbf{X}^j \in R^{5\times3\times(2r+1)\times(2r+1)} \rightarrow$ $2\times$(3DConv$\rightarrow$ReLU$\rightarrow$3DBN)$\rightarrow$ $\mathbf{Z}_x^j \in R^{c_1,h,w}$

$\mathbf{S}^j$ is passed to a 2D Convolutional branch that "convolve" with the input along the spatial domain in two sequential 2D convolutional layers (conv. layers) and return a 3D tensor of high level features, $\mathbf{Z}_s^j$. We set the model to have convolutional filters in both branches of the same spatial size so that both 3D tensors of high-level filters evolving from the two branches to have the spatial size,$(h\ timesw)$. We summarie this branch as follows: $\mathbf{S}^j \in R^{2\times(2r+1)\times(2r+1)} \rightarrow 2\times$(2DConv$\rightarrow$ReLU$\rightarrow$2DBN)$\rightarrow$ $\mathbf{Z}_s^j \in R^{c_1,h,w}$

This two 3D tensors of high-level features,$\mathbf{Z}_x^j, \mathbf{Z}_s^j$, returned form each branch are then stacked along their third domain to form the 3D tensor $\mathbf{Z}^j$, and propagated it to the rest of the network. The final part of the network has another two 2D convolutional layers and after the last convolutional operation, the output is propagated to a SPP layer, two FC layers with DO in between and a sigmoid squashing function as in our CNN model. We summarise the final part of the network as follows: $\mathbf{Z}^j \in R^{(c_1+c_2),h,w} \to 2\times$(2DConv→ReLU→2DBN) → SPP($n$, **k**) → FC(spp,100) → ReLU → 1DBN → DO(p) → FC(100,1) → $\sigma \to \hat{Y}^j_{t+1}$

In this model architecture, again, we utilized 2DBN and 3DBN between each 2D and 3D conv. layers after applying ReLU activation function. The number of filters in each of the 2D and 3D conv layers was set as free parameter, as well as the spatial size of the filters. Parameters of SPP and DO layers are allowed to vary too. The model is able to analyze tensors of any spatial size.This model flexibility allowed us to experiment with its architecture. Results are shown in Chapter 7. Figure S11 illustrate our second model architecture.

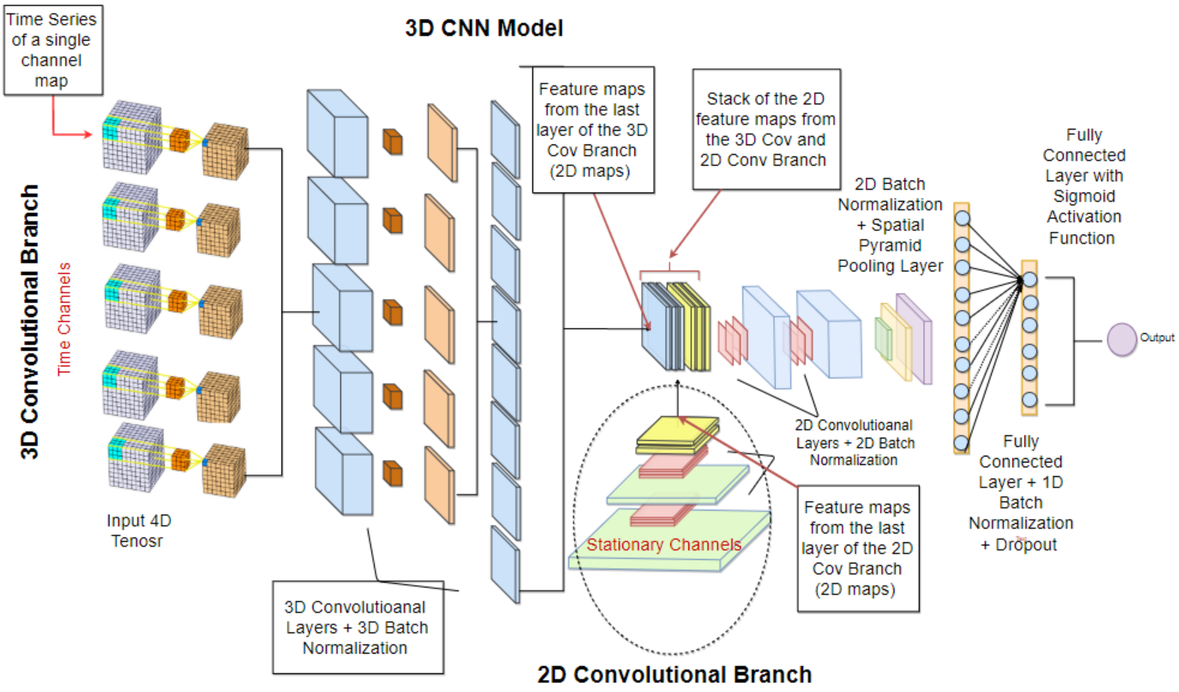

**Figure S11: Model 3:** 3D CNN Model. Architecture summary:
$\mathbf{X}^j \in R^{5\times3\times(2r+1)\times(2r+1)} \to 2\times$(3DConv→ReLU→3DBN)→ $\mathbf{Z}^j_x \in R^{c_1,h,w}$
$\mathbf{S}^j \in R^{2\times(2r+1)\times(2r+1)} \to 2\times$(2DConv→ReLU→2DBN)→ $\mathbf{Z}^j_s \in R^{c_2,h,w}$
$\mathbf{Z}^j \in R^{(c_1+c_2),h,w} \to 2\times$(2DConv→ReLU→2DBN) → SPP($n$, **k**) → FC(spp,100) → ReLU → 1DBN → DO(p) → FC(100,1) → $\sigma \to \hat{Y}^j_{t+1}$

#### S3.4 Model Architectures 3 & 4: Convolutional and Deep Convolutional Long Short Term Memory Recurrent Neural Network

While our second, 3D CNN, model was able to utilize all three dimensions of the input data, it has feed-forward learning. As we discussed in Chapter 2.3, when one has sequential data, recurrent learning is usually very powerful learning process. This inspire us to design our next two models so that they learn the temporal structure of the input data recurrently. More precisely, they both have Convolutional Long Short Term Memory cell, or cells, in their architecture (see Chapter 2.3.1 for more details about the cell structure).

Both our Model 3 and Model 4 have architecture similar to that of our 3D CNN model. For a data pint $j$, they take as an input its static tensor $\mathbf{S}^j \in R^{2\times(2r+1)\times(2r+1)}$ and its time series $\{\mathbf{X}^j_{t-2}, \mathbf{X}^j_{t-1}, \mathbf{X}^j_t\}$, each $\in R^{5\times(2r+1)\times(2r+1)}$. Again the static tensor $\mathbf{S}^j$ and the time series of tensors $\{\mathbf{X}^j_k\}^t_{k=t-3}$ are separately propagated through two separate branches, **(Enocoder + Conv LSTM RNN)** and **2D CNN Static**, each of which return 3D tensors of high-level features ,$\mathbf{Z}^j_x, \mathbf{Z}^j_s$. $\mathbf{Z}^j_x, \mathbf{Z}^j_s$ are then stacked together along their channel axis to form $\mathbf{Z}^j$. This joint 3D tensor is propagated through the rest of the network. Finally, the model return $\hat{Y}^j_{t+1}$ where $j \in \mathbf{J}_t$ (see Chapter 4 for more details about this notation).

Here we give detailed explanation what these two branches does. The branch that takes $\{\mathbf{X}^j_{t-2}, \mathbf{X}^j_{t-1}, \mathbf{X}^j_t\}$ of both Model 3 and Model 4 differs form this in our second model,3D CNN, in two ways, it has Encoder sub-branch and a Convolutional Long Short Term sub-branch, **ConvLSTM**.

The **Enocoder** encodes the time series of 3D tensors $\{\mathbf{X}^j_k\}^t_{k=t-3}$ to another time series of 3D high-level features $\{\tilde{\mathbf{X}}^j_k\}^t_{k=t-3}$. It does so by feeding each $X^j_{t_j}$ through the same three 2D conv.layers. We summarise this sub-branch as follows:$\mathbf{Enocoder}:$
$\{\mathbf{X}^j_k\}^t_{k=t-3} \rightarrow \{\tilde{\mathbf{X}}^j_k\}^t_{k=t-3}:$
$\mathbf{X}_t \in R^{5\times(2r+1)\times(2r+1)} \rightarrow 3\times$(2DConv→ReLU→2DBN) $\rightarrow \tilde{\mathbf{X}}_\mathbf{t} \in R^{c_1\times h_1\times w_1}$
$\mathbf{X}_{t-1} \in R^{5\times(2r+1)\times(2r+1)} \rightarrow 3\times$(2DConv→ReLU→2DBN) $\rightarrow \tilde{X}_{t-1} \in R^{c_1\times h_1\times w_1}$
$\mathbf{X}_{t-2} \in R^{5\times(2r+1)\times(2r+1)} \rightarrow 3\times$(2DConv→ReLU→2DBN) $\rightarrow \tilde{\mathbf{X}}_{\mathbf{t-2}} \in R^{c_1\times h_1\times w_1}$

This time series of high-level 3D features, $\{\tilde{\mathbf{X}}^j_k\}^t_{k=t-3}$, is then handled recurrently buy the **ConvLSTM** sub-branch.**ConvLSTM** of Model 3 pass this new time series to one 2D Convolutional Long Short Term Memory cell and after the last iteration of this cell, this branch of the model returns a 3D feature tensor, $\mathbf{Z^j_x}$, that stores the essential spatio-temporal "memory" of the model input. Model 4 differs from Model 3 in that its **ConvLSTM** sub-branch is "deep". More precisely, the time series of high-level features, $\{\tilde{\mathbf{X}}^j_k\}^t_{k=t-3}$, is passed to a stack of several, say $d$, ConvLSTM cells. The latest in time output of the deepest cell is returned as a 3D feature tensor that stores the essential spatio-temporal "memory", $\mathbf{Z^j_x}$. We sumarise this sub-branch of Model 4 as follows: **Conv LSTM RNN:** $\{\tilde{\mathbf{X}}^j_k\}^t_{k=t-3} \rightarrow d\times$(ConvLSTM) $\rightarrow \mathbf{Z}^j_x \in R^{c_2,h_2,w_2}$

The branch of Model 3 and 4 taking $\mathbf{S}^j$ is identical and also very similar to that of Model 2. $\mathbf{S}^j$ is passed to three sequential 2D convolutional layers (conv. layers) and return a 3D tensor of high level features, $\mathbf{Z}^j_s$. Here again we set the model to have convolutional filters in this branch such that the output of this branch has the same spatial size as the output of the **ConvLSTM** sub-branch. Thus, this branch, which we denote as **2D CNN Static**, returns $\mathbf{Z}^j_s \in R^{c_3,h_2,w_2}$. We summarise this branch as follows: **2D CNN Static:** $\mathbf{S}^j \in R^{2\times(2r+1)\times(2r+1)} \rightarrow 3\times$(2DConv→ReLU→2DBN)→

$\mathbf{Z}_s^j \in R^{c_3,h_2,w_2}$

Finally, the outputs of the two branches,$\mathbf{Z}_x^j$,$\mathbf{Z}_s^j$, are stacked together along the channel axis to form $\mathbf{Z}^j \in R^{(c_2+c_3),h_2,w_2}$ which is then propagated through the rest of the network. We denote this part of the network as **2D CNN Joint**. It consists of three 2D conv. layers, SPP layer and two 2 FC layers with DO layer in between. The output of the last FC layer is squashed with sigmoid non-linearity, $\sigma$, and has value $\hat{Y}_{t+1}^j \in [0,1]$.We summarise this propagation as follows: **2D CNN Joint** $\mathbf{Z}^j \in R^{(c_2+c_3),h_2,w_2} \rightarrow$ 3×(2DConv→ReLU→2DBN) $\rightarrow$ SPP($n$, $\mathbf{k}$) $\rightarrow$ FC(spp,100) $\rightarrow$ ReLU $\rightarrow$ 1DBN $\rightarrow$ DO(p) $\rightarrow$ FC(100,1) $\rightarrow \sigma \rightarrow \hat{Y}_{t+1}$

Figure S12 illustrate the model architecture of Model 4, that has $d = 2$ ConvLSTM cells one above another in its **ConvLSTM** sub-branch. Here again, we have designed our models to be flexible with respect to their hyper-parameters. For Model 4 one can explore how the model performance change when the number of filters and their spatial size of each layer present in the network take different values, by changing the depth, $d$, of the ConvLSTM network section, by controlling the overfitting effect with different dropout ratio, p, and the downsampling rate of the parameters of the SPP layer. Finally, as with our other models, the input tensors can be of any spatial size $((2r + 1) \times (2r + 1))$.

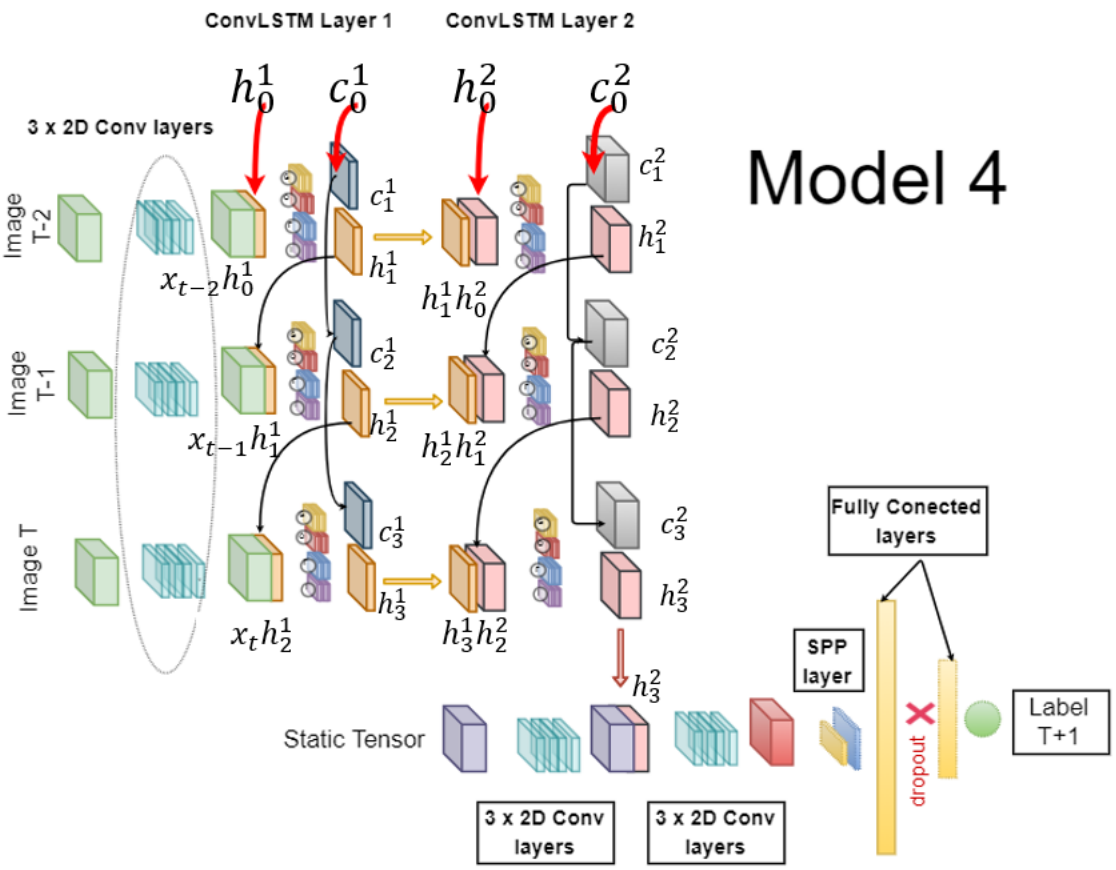

**Figure S12:** Model 4: Deep ConvLSTM Neural Network with 2 ConvLSTM cells. Architecture summary:

**Enocoder:** $\{\mathbf{X}_k^j\}_{k=t-3}^t \rightarrow \{\tilde{\mathbf{X}}_k^j\}_{k=t-3}^t$

$\mathbf{X}_t^j \in R^{5\times(2r+1)\times(2r+1)} \rightarrow 3\times$(2DConv$\rightarrow$ReLU$\rightarrow$2DBN) $\rightarrow \tilde{\mathbf{X}}_t^j \in R^{c_1\times h_1\times w_1}$

$\mathbf{X}_{t-1}^j \in R^{5\times(2r+1)\times(2r+1)} \rightarrow 3\times$(2DConv$\rightarrow$ReLU$\rightarrow$2DBN) $\rightarrow \tilde{\mathbf{X}}_{t-1}^j \in R^{c_1\times h_1\times w_1}$

$\mathbf{X}_{t-2}^j \in R^{5\times(2r+1)\times(2r+1)} \rightarrow 3\times$(2DConv$\rightarrow$ReLU$\rightarrow$2DBN) $\rightarrow \tilde{\mathbf{X}}_{\mathbf{t-2}}^{\mathbf{j}} \in R^{c_1\times h_1\times w_1}$

**Conv LSTM RNN:** $\{\tilde{\mathbf{X}}_k^j\}_{k=t-3}^t \rightarrow 2\times$(ConvLSTM) $\rightarrow \mathbf{Z}_x^j \in R^{c_2,h_2,w_2}$

**2D CNN Static:**

$\mathbf{S}^j \in R^{2\times(2r+1)\times(2r+1)} \rightarrow 3\times$(2DConv$\rightarrow$ReLU$\rightarrow$2DBN)$\rightarrow \mathbf{Z}_s^j \in R^{c_3,h_2,w_2}$

**2D CNN Joint:**

$\mathbf{Z}^j \in R^{(c_2+c_3),h_2,w_2} \rightarrow 3\times$(2DConv$\rightarrow$ReLU$\rightarrow$2DBN) $\rightarrow$ SPP($n$, k) $\rightarrow$ FC(spp,100) $\rightarrow$ ReLU $\rightarrow$ 1DBN $\rightarrow$ DO(p) $\rightarrow$ FC(100,1) $\rightarrow \sigma \rightarrow \hat{Y}_{t+1}^j$

### S4 Methodological notes

Data within the training periods was split into training, validation and test sets by 3:1:1.

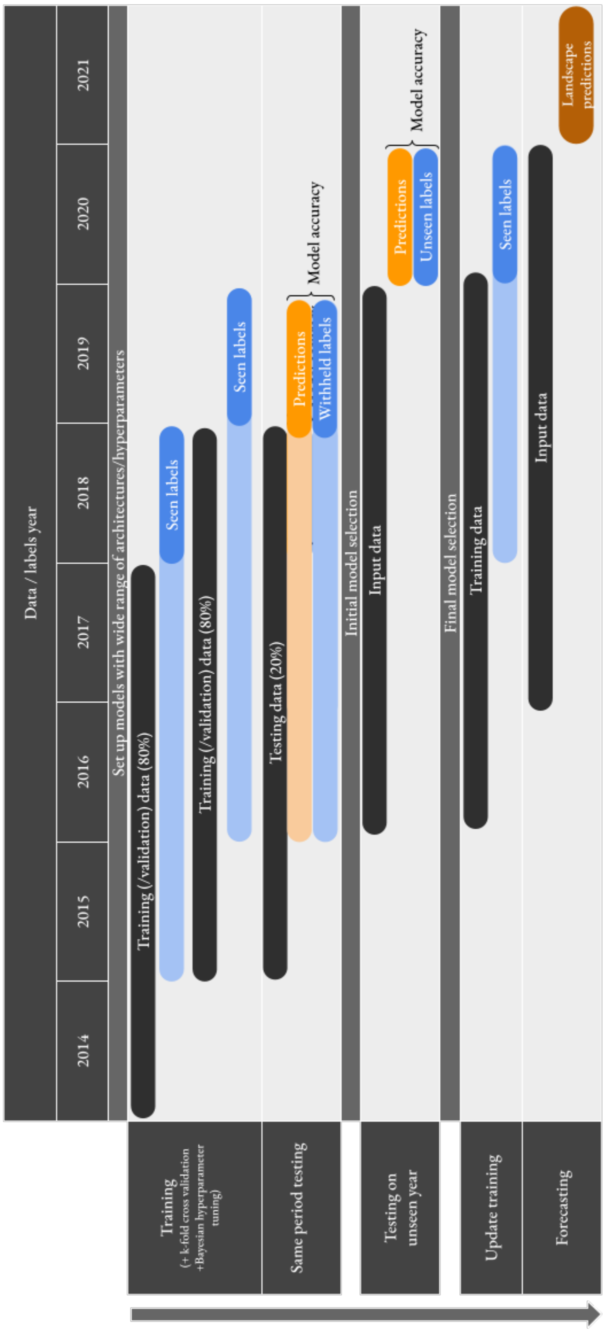

**Figure S13:** Arrangement of training, testing and forecasting processes

#### S4.1 Training

For Model Architectures 2, 3 and 4 models were first trained on 2014-2017 input data matched with 2018 labels. They were then trained on 2015-2018 input data on 2019 labels.

Model Architecture 1 was trained on 2014 input with 2015 labels, 2015 input with 2016 labels, 2016 input with 2017 labels, 2017 input with 2018 and 2018 input with 2019 labels all at once.

5-fold cross validation took place during the training routines to track how well the models were learning.

#### S4.2 Model tuning

For the initial broad model inter-comparison we conducted a grid search to find effective model parameters. For the development of the most promising models we used a Bayesian hyperparameter sweep Snoek et al. (2012) as implemented on WandB.

#### S4.3 Testing

To understand how well models would forecast they had to be tested on a time period that was outside of the training period. To do this we see how well they can predict deforestation in 2020 (an unseen year for the model). The models that can predict deforestation that happened 2020 with the best accuracy are identified and their parameters are recorded.

#### S4.4 Forecasting

The models with the best parameters were updated with training data from 2015 to 2019 (with 2020 labels). The model weights were retained to keep the learning that had taken place on the 2014-2018 data.

The updated model is used to forecast deforestation for 2021. It produced a risk map across all forested land in the area of interest. A threshold based on anticipated overall levels of deforestation could be applied to the map at a required level (e.g. from 0 to 1) to identify pixels that are likely to undergo deforestation.

#### S4.5 Class Imbalance Problem

The number of pixels that become deforested in Madre de Dios 2017 and 2018 accounted for around 0.30%, 0.26% of the total number of pixels covering the forested study area on which our models were trained. This ratio of 99.7 : 0.3 made our dataset extremely imbalanced. Therefore, by simply prediction all pixels as forested one would get misleading accuracy of 99.7%, but such model will useless model. To address this issue we took several steps. Our objective was to be able to identify the top 20% most susceptible areas with high accuracy so that attention could be. In reality, a perfect model predicting up to 20% deforestation labels would be able to achieve no more than 80.3% accuracy as 19.7% non-deforested pixels would be labelled as suspected. Therefore, when testing and training our models we needed to make class-aware sampling. Recently, this class imbalance problem was addressed by Buda et al. (2018) in the context of Deep Learning models. They examined the impact such an imbalance can have on CNNs classifiers via several experiments and empirically concluded that the effect of it is detrimental. They also investigated the efficiency of several algorithms dealing with class imbalance, commonly used in the Machine Learning field, when applied on Deep Learning methods. Their results shows that the best approaches in the DL field are oversampling, or if training time is a problem, undersampling is an alternative, where sampling must be done to the extend where the imbalance is completely removed.

While we assumed (Buda et al., 2018) conclusion that undersampling is the right approach when dealing with imbalanced data, the extend to which this should be done we set as a "free",tuning, parameter. Our reasoning for this is that the cost of missing to detect future deforestration event is much higher than raising a wrong deforestation alarm.

To explore what should be the ratio of forested to deforested pixels in our train, validation and test data, we developed a**sampling algorithm**, which when given a dataset $\mathbf{D}_t$ and a parameter $\theta$, returns a new undersampled dataset $\tilde{\mathbf{D}}_t$. This new dataset $\tilde{\mathbf{D}}_t$ has all data-points of the the minority class, pixels labeled as deforested $(Y^j_{t+1} = 1,\ j \in \mathbf{J}_t)$, but the number of observations form the majority class, forested pixels$(Y_{t+1} = 0\ j \in \mathbf{J}_t)$, is set to be such that the ratio of the two calses in $\tilde{\mathbf{D}}$ is $\theta$. The observations from the majority class are randomly selected and at each call of this algorithm, the set of observation drawn from the majority class was updated with new, distinct, set of randomly selected forested observations. The following section explains how we utilized this algorithm in our models training process.

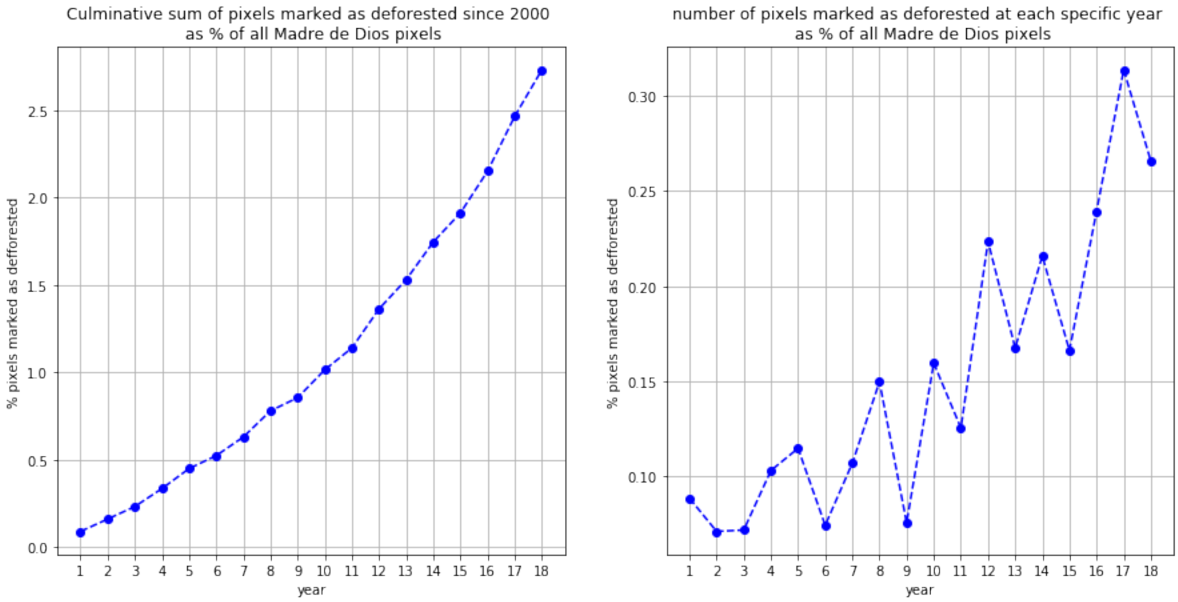

**Figure S14:** Percentage deforested pixels of all pixels covering Madre De Dios area

#### S4.6 Early Stopping regularization

In our training process we used Adam optimizer (Kingma, Ba, 2014) and Binary Cross Entropy Loss.

Early stopping is a regularization technique that is used when there is an iterative learning process. Several variations of this technique exist, but the one we utilize aims to make the model a better fit to data outside of the training set, validation dataset.

Here we describe the way we employed early stopping regularization. After each epoch, the model under training was set to interference state and its performance was evaluated on a separate validation data set. The ratio of training data size to validation data size was kept to 60:20. Each data set has equal ratio of 0 and 1 labels, the ratio of forested to deforested pixels labels, which was set to $\theta$. At each epoch iteration, if the performance of the model on this validation data set was better than its performance in the previous iteration according to a certain criterion, the leering process continue to iterate through the batches one more time until new epoch is completed. If however, the performance on this iteration is worse, a counter variable starts counting how many times the updated model after each epoch is worse than the previously best one. That is, if the model gives tree times in a row, i+1, i+2, i+3, worse performance than the one recorded at iterations i, the counter has value 3. The counter counts up to pre-defined “waiting time” and is reset to 0 each time a new, better performing model is found. In our training strategy we set the “weighting time” to be 3. Which means that if our model performs three times in a row worse than it did at iteration i, then we stop the learning process, and chose the model with weights evaluated at iteration i. If the model however performs worse at iteration i+1 and i+2 than it did at i, but at i+3 it is better according to the criteria, the counting process restart and the model. The model at iteration i+4 is now compared with the previously best, i+3.

The above described algorithm is general, and well known within DL community. The way we modified this algorithm is by re-sampling our taring and validation data sets after each epoch by calling our sampling algorithm ( as described in the previous Section S4.5) with input parameter , which preserves the ratio of 0 to 1 labels in both train and validation data set, but update the under-sampled class of 0 labels, with new, distinct, randomly sampled 0 labels. We also investigated the effect of employing this Early Stopping technique with different criteria, namely the Area Under The Receiver Operating Characteristics (AUC), Weighted Binary Cross entropy Loss and the Cost from Cost matrix. As training time was an issue, we only run our algorithm up to 5 epochs, with "waiting time" 3 and the best performing model was returned. Figure **??** illustrate the training process of three different models, where for each different criterion was selected and demonstrate the advantages of utilizing Early Stopping regularization.

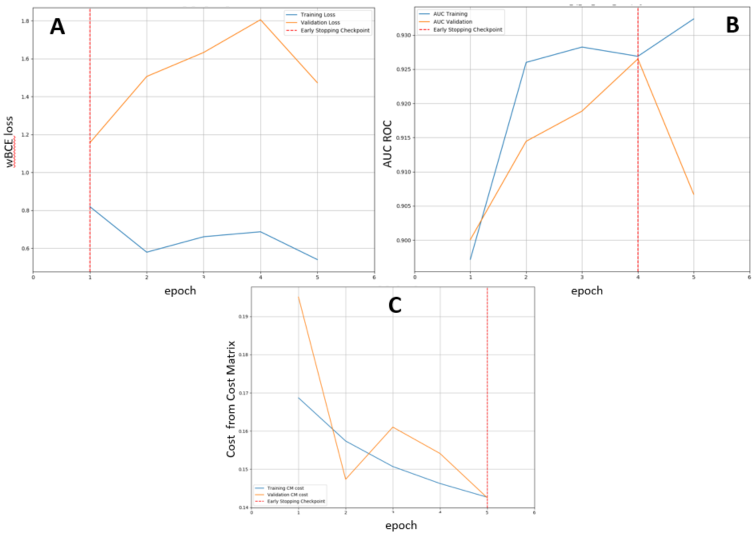

**Figure S15:** Training with an Early Stopping regularization.
**A** The selected criterion is Weighted Binary Cross entropy Loss. The selected model is the one learning its weights up to epoch 1. The model overfits onwards. While the loss evaluated on the train data decrease (blue line), on validation data the loss it increase (yellow line) with minimum validation loss recorded at epoch 1.
**B** The selected criterion is AUC ROC. The selected model is the one learning its weights up to epoch 4. The model does not overfit up to epoch 4. However, the AUC on validation data decrease at the next epoch 5. The learning process is ended due to time limitations.
**C** The selected criterion is Cost from Cost matrix. The selected model is the one learning its weights up to last epoch 5 when both train and validation cost was at its lowest figure. The can not be said to overfit. However, while a smooth decrease of the cost was observed on train data, the validation cost has drop at epoch 2 and spike at epoch three.

#### S4.7 Training models with mono-temporal and multi-temporal inputs

Our four models ability to correctly forecast deforestation event is compared on labels corresponding to year 2018, $\hat{Y}^j_{2018}, j \in \mathbf{J}_{2017}$. The input data for our last three models, Model 2, Model 3, Model 4, is $(S^j, X^j_{2015}, X^j_{2016}, X^j_{2017)|j \in \mathbf{J}_{2017}}$ whereas Model 1 takes as an input $(S^j, X^j_{2017)|j \in \mathbf{J}_{2017}}$.

When training, the only possible training data for Model 2, Model 3, Model 4 is $(S^j, X^j_{2014}, X^j_{2015}, X^j_{2016)|j \in \mathbf{J}_{2016}}$ with corresponding labels $\hat{Y}^j_{2017}, j \in \mathbf{J}_{2016}$. This data, before being undersampled (with ratio of 0:1 labels = 99.7 : 0.03), was divided into training, validation and test data subsets with ratio 6:2:2. The taring and validation data sets were use for training with an Early Stopping regularization. We used the test data to select the best "tuned" model of each class. The ratio of 0 to 1 labels was kept the same in all three subsets, training, validation and test. Sampling algorithm (Section S4.5) with parameter  was then used on each of the data subsets when we examined the effect of different imbalance ratios.However, when training our CNN model, different time set ups for training data are possible. In the subsection bellow we discuss how we addressed that issue.

#### S4.8 Training and testing CNN models with different time set up

Our first, 2D CNN, model takes a singe 3D-tensor as an input $\mathbf{SX}^j_t \in R^{7\times 2r+1\times 2r+1}$ and output a integer score $\hat{x}^j_t \in [0,1]$ that can be interpreted as our confidence level of pixel $j$ to be deforested in year t+1. Then, for specified threshold $tr$ we assign $Y^j_{t+1} = I_{x_t > tr}$. Considering the above notations, the data set for a CNN model is then defined as $D_t = \{(\mathbf{SX}^j_t], Y^j_{t+1})|j \in \mathbf{J}_t\}$ if only one year data is used. However, one can also use the more than one year data by disregarding the time index of the data but only using features with one year lag from their predicted labels. To be more precise, a set up as $\mathbf{D}_{t_k} = D_{t-k} \cup D_{t-(k-1)} \cup .... \cup D_{t-1} \cup D_t$ is possible, in which case k+1 years of data is utilized. This possibilities made us ask several questions. Firstly, what is the optimal number of year, k? Secondly, if one year is used, is our first model time invariant? That is, if we train it on $D_{2015} = \{(\mathbf{SX}^j_{2015}], Y^j_{2016})|j \in \mathbf{J}_{2015}\}$, how it performs on the two data-sets $D_{16} = \{(\mathbf{SX}^j_{16}], Y^j_{17})|j \in \mathbf{J}_{2016}\}$ and $D_{2015} = \{(\mathbf{SX}^j_{2017}], Y^j_{2018})|j \in \mathbf{J}_{2017}\}$?. Finally, to be compared with the rest of our models, that have taring and validation and test data sets with labels in 2017, how should we train our CNN model so as to be equally biased towards labels in 2017 while simultaneously utilizing all years data.

To answer the last, we did as follows, while the training data year was allowed to vary. However, for validation and test data we used 20% and 20% of the original $D_{2016}$ data (where the ratio of 0:1 labels was 99.7 : 0.03) respectively. The rest 60% were considered as part of the training data. We then applied Sampling algorithm(Section S4.5) with parameter on each of the data subsets to examine the effect of different imbalance ratios .

After we selected our best 2D CNN model, we used its hyper parameters to train a model on $D_{2015} = \{(\mathbf{SX}^j_{2015}], Y^j_{2016})|j \in \mathbf{J}_{2015}\}$ and examined how it performs on the two data-sets $D_{2016} = \{(\mathbf{SX}^j_{2016}], Y^j_{2017})|j \in \mathbf{J}_{2016}\}$ and $D_{2017} = \{(\mathbf{SX}^j_{2017}], Y^j_{2018})|j \in \mathbf{J}_{2017}\}$.

### S5 Computation

#### S5.1 Software

The development of the neural networks and processes to train and test them were done using the PyTorch library Version 1.9.0.

An Anaconda virtual environment was used to manage package dependencies. The environment to perform all tasks relevant to this project (*forecast*) is saved on the project GitHub repository as a *.yml* file.

#### S5.2 Computing resources / hardware

A High Performance Computing cluster was used to train, test and forecast. Typically, tasks ran with four Tesla P100-PCIE-16GB GPUs and twelve 5980MB CPUs up to twelve hours although this could vary depending on resource availability and task requirements.

### S6 Experimental set up

#### S6.1 Model 1: 2D CNN model

LINK TO SCRIPTS

To “tune” our first model we selected a base of hyper parameters that we found works well by guided guess. We then changed each of them at a time and recorded how the model will perform on a test data with labels in 2017.

Our base model was trained on the full set of data available,

$$\mathbf{D}_{2014-2016} = D_{2014} \cup D_{2015} \cup D_{2016} =$$

$$\{(\mathbf{SX}^j_{2014}], Y^j_{2015})|j \in \mathbf{J}_{2014}\} \cup \{(\mathbf{SX}^j_{2015}], Y^j_{2016})|j \in \mathbf{J}_{2015}\} \cup (\mathbf{SX}^j_{2016}], Y^j_{2017})|j \in \mathbf{J}_{2016}\}$$

However, in order to make it equally biased towards labels in year 2018 as the rest of our models are, we only only used 60% of $(\mathbf{SX}^j_{2016}], Y^j_{2017})|j \in \mathbf{J}_{2016}\}$ for training and devoted the rest 20% and 20% for validation and test data [UPDATE TO DESCRIBE K-FOLD CROSS VALIDATION]. When splitting the data, no undersampling was performed and the split was done so that the original ratio of 0 to 1 labels were preserved in all three data subsets (99.688 : 0.312). Thus 64803129, 21601044 and 21601046 pixels were allocated for the train, validation and test data sets respectively with the total size of the 2016 data being $|\mathbf{J}_{2016}|$ = 108005219. For train and validation undersampling of our base model we used $\theta = 1$, or said differently, the ratio of 0 to 1 labels was modified to 1:1. Thus, after undersampling, the training and validation data sizes were decreased to 404978 and 134994 respectively. Unless explicitly said in the model experiment set up (case 12), after each epoch we changed the training data set with one year older data, that was again undersampled to have ratio $\theta = 1$. Thus, at epoch 1, the 60% of the undersampled 2016 data ,$\tilde{D}^{60\%}_{2016}$, was used for training and the 20% ,$\tilde{D}^{20\%}_{2016}$, allocated for validation. At epoch 2, new undersampled data was added for training , namely $\tilde{D}_{2015}$ , whereas the validation data was re-sampled with new 2016 zero values via our **Sapling algorithm** S4.5 with $\theta$ again = 1. For epoch 3 the same procedure was performed with new training data being $\tilde{D}_{2014}$. If the model was allowed to further iterate to epoch 4 and 5, the training data was newly undersampled $\tilde{D}^{20\%}_{2016}$ and $\tilde{D}_{2015}$.

The default Early Stopping Criterion was AUC, the weighting time 3 and the maximum number of epoch iterations 5. Weighted BCE with positive weights 5 was used as loss to be optimize. We used Adam optimizer (Kingma, Ba, 2014) with default learning rate of 0.0001. The batch size of all training experiments in this report was set to 80.

The default model architecture was as follows:

Input tensor $\mathbf{SX}^j_t \in R^{7\times 35\times 35} \rightarrow 4\times$(2DConv$\rightarrow$ReLU$\rightarrow$2DBN) $\rightarrow$ SPP$(2, [13, 5])$ $\rightarrow$ FC(1552,100) $\rightarrow$ ReLU $\rightarrow$ 1DBN $\rightarrow$ DO(0.2) $\rightarrow$ FC(100,1) $\rightarrow \sigma \rightarrow \hat{Y}^j_{t+1}$

That is, the default input spatial dimensions was (35×35) and it was propagated to 4 2D conv. layers that decreased its spatial size form 35 to 25. Conv. layer 1 had 8 filters each with spatial size (5×5), conv. layers 2 and 3 had 16 filters each with spatial size (3×3),and finally conv. layer 4 had 8 filters each with spatial size (3×3). All slided at stride 1 with no padding applied. Then a SPP layer downsampled it to a 1d vector of size 1552 via 2 max pooling filters returning 3D tensors of spatial size $13 \times 13$ and $5 \times 5$ respectively. This was fed to the two fully conceded layers which had dropot ratio 0.2. We then let examined how the network will perform if the the following hyperparameters were changed: shown: dropout ratio: 0.5, dropout ratio: 0, $\theta : |Y = 0| : |Y = 1|$ = 1:2 = 0.5, Input spatial size: $45 \times 45$, SPP with n = 2 and $\mathbf{k} = [15, 5]$, Number of filters in conv. layers 1 to 4: 16, Early Stopping criterion: BCE loss, BCE loss as loss to be optimized, Weighted BCE loss with positive weight 2 as loss to be optimized, Learning rate of Adam: 0.001, use only year $D_{2016}$, Results of these experiments are shown in Table **??**

#### S6.2 Model 2: 3D CNN model

LINK TO SCRIPT

For all other three models the full time series of data was utilized:

$$\{(S^j, X^j_{2014}, X^j_{2015}, X^j_{2016}) | j \in \mathbf{J}_{2016}\}$$

It was split to train, validation and test data sets with ratio 6:2:2 in such a manner that the original ratio of 0 to 1 labels was preserved (99.69 : 0.31). Thus 64803129, 21601044 and 21601046 pixels were allocated for the train, validation and test data sets respectively with the total size of the 2016 data being $|\mathbf{J}_{2016}|$ = 108005219. For train and validation undersampling of our base model we used $\theta = 1$, or said differently, the ratio of 0 to 1 labels was modified to 1:1. Thus, after undersampling, the training and validation data sizes were decreased to 404978 and 134994 respectively.

Here again for default training set up we used weighted BCE loss with positive weight 3, ADAM optimizer with learning rate 0.0001 and used AUC as an early stopping criterion.

The model architecture was as follows:
Branch 1 : $\mathbf{X}^j \in R^{5\times3\times35\times35} \to 2\times$(3DConv→ReLU→3DBN)→ $\mathbf{Z}^j_x \in R^{c_1,h,w}$
Branch 2: $\mathbf{S}^j \in R^{2\times35\times35} \to 2\times$(2DConv→ReLU→2DBN)→ $\mathbf{Z}^j_s \in R^{c_2,h,w}$
Branch 3: $\mathbf{Z}^j \in R^{(c_1+c_2),h,w} \to 2\times$(2DConv→ReLU→2DBN) → SPP$(1, 15)$ → FC(spp,100) → ReLU → 1DBN → DO(0.3) → FC(100,1) → $\sigma \to \hat{Y}^j_{t+1}$

That is, each of the 2 3D conv. layers of the branch taking $\mathbf{X}^j \in R^{5\times3\times35\times35}$ have 16 4D filters of dimension $R^{5,2,3,3}$ which slide at stride 1 and no padding was applied. Each of the 2 2D conv. layers of the branch taking $\mathbf{S}^j \in R^{2\times35\times35}$ have 8 3D filters of of dimension $R^{2,3,3}$ which slide at stride 1 and no padding is applied. Finally, the branch taking $\mathbf{Z}^j \in R^{32,31,31}$ has 4 3D filters of dimension $R^{32,3,3}$ which slide at stride 1 in each of its 2 2D conv. layers. No padding is applied. The SPP layer has one max pooling filter that returns a 3D tensor of dimension $R^{4,13,13}$ which is flattened and passed to the first fully connected layer. A dropout ratio before the final FC layer of rate 0.3 is applied.

We tested this base set up against: dropout = 0.5, larger spatial sizes of the filters in the first two brances - filters of branch 1 and branch 2 $\in R^{5,2,5,5}$ and $\in R^{2,5,5}$ respectively, increased number of filters in each of the three branches - 8, 32, 8 respectively, SPP with n=2 and $\mathbf{k} = [12, 5]$ , Cost of Confusion Matrix as an early stopping criterion, input image size 31 and 45, $\theta = 0.5$, weighted BCE loss with weight 10, L2 regularization with parameter 0.6 and finally, learnnig rate of ADAM 0.001. Table S4 summarize the results of this experiments.

#### S6.3 Model 3 and 4: ConvLSTM RNN model Deep ConvLSTM RNN model

For our last two models, Model 3: ConvLSTM RNN and Model 4: Deep ConvLSTM RNN, the trainig, validation and test data set up was identical as this of Model 2. Base Model 3 differed form base Model 4 only in that Model 4 has two identical ConvLSTM cells one after another, which defined it as "deep".

The base set up was weighted BCE loss with positive weight w = 5, AUC as an early stopping criterion, Adam optimizer with learning rate 0.0001, $\theta = 1$.

The base models architecture is as follows:
**Enocoder:**
$\{\mathbf{X}_k^j\}_{k=t-3}^t \in R^{5\times 25\times 25} \rightarrow 3\times$(2DConv$\rightarrow$ReLU$\rightarrow$2DBN) $\rightarrow \{\tilde{\mathbf{X}}_k^j\}_{k=t-3}^t \in R^{c_1\times h_1\times w_1}$
**Conv LSTM RNN:** $\{\tilde{\mathbf{X}}_k^j\}_{k=t-3}^t \rightarrow 2\times$(ConvLSTM) $\rightarrow \mathbf{Z}_x^j \in R^{c_2,h_2,w_2}$
**2D CNN Static:**
$\mathbf{S}^j \in R^{2\times 25\times 25} \rightarrow 3\times$(2DConv$\rightarrow$ReLU$\rightarrow$2DBN)$\rightarrow \mathbf{Z}_s^j \in R^{c_3,h_2,w_2}$
**2D CNN Joint:**
$\mathbf{Z}^j \in R^{(c_2+c_3),h_2,w_2} \rightarrow 3\times$(2DConv$\rightarrow$ReLU$\rightarrow$2DBN) $\rightarrow$ SPP$(1,12)$ $\rightarrow$ FC(spp,100) $\rightarrow$ ReLU $\rightarrow$ 1DBN $\rightarrow$ DO(0.3) $\rightarrow$ FC(100,1) $\rightarrow \sigma \rightarrow \hat{Y}_{t+1}^j$

That is, the input spatial size is 25x25. The **Enocoder** branch have 3 2D conv. layers with 8 filters each, of dimension $\mathbf{R}^{5,3,3}$ that slides at stride 1 and no padding is applied.

The **Conv LSTM RNN** than take $\{\tilde{\mathbf{X}}_k^j\}_{k=t-3}^t \in R^{8\times 19\times 19}$ pass it to $2\times$(ConvLSTM) ($1\times$(ConvLSTM) for Model 3) and return $\mathbf{Z}_x^j \in R^{8,19,19}$. Both ConvLSTM cells have 8 filters for each of the gates, with each filter $\in R^{8,3,3}$. However, a padding of size 1 is applied in order the cell to be recurrent. Each filter slides at stride 1.

Both **2D CNN Joint** and **2D CNN Static** have 8 filters in each of their 3 2D conv. layers, each of spatial dimension 3x3, sliding at stride 1 and no padding is applied. The SSP layer has one maxpooling filter that returns 3D tensor $\in R^{8,12,12}$. Finally, dropout of rate 0.3 is applied between the two FC layers.

We compared these models, against this that has the following changes: dropout = 0.5, for Model 3 the number of filters in each of the for branches was changed from (8,8,8,8) to (8,8,4,4), (16,8,4,4), (16,16,4,4), and for Model 4 from (8,(8,8),8,8) to (16,(8,8),8,8), (16,(16,16),8,8), (8,(8,8),4,4), Where the first input of the notation (.,.,.,.) is the numer of filters in the **Enocoder**, the second for $2\times$(ConvLSTM) / $1\times$(ConvLSTM) for Model 3 and so on. We also compared models when for early stopping criterion the Confusion Matrix was evaluated, when the input have size 31, 35, wen $\theta = 0.5$, L2 regularization with parameter 0.2 and finally, ADAM with learning rate 0.001. The results of this expreiments are dipsplayed in Tables S5 and S6 for Model 3 and 4 respectively.

### S7 Experimental Results

#### S7.1 Broad model intercomparison (Madre de Dios)

##### S7.1.1 Model performance within training period

**Table S3:** Model 1, Experimental results

| hyperparameter | epochs | Total time | AUC on Train | AUC on Valid. | AUC on Test |
|---|---|---|---|---|---|
| base | 3 | 3.2h | 0.924 | 0.921 | 0.921 |
| dropout ratio: 0.5 | 5 | 6h | 0.935 | 0.918 | 0.917 |
| dropout ratio: 0 | 1 | 2.5h | 0.911 | 0.922 | 0.921 |
| $\theta$ : 0.5 | 4 | 1.7h | 0.930 | 0.934 | 0.933 |
| In size: $45 \times 45$ | 4 | 5h | 0.943 | 0.950 | **0.951** |
| SPP: n = 2,k = $[15, 5]$ | 4 | 4h | 0.936 | 0.918 | 0.919 |
| hidden dim = 16 | 4 | 4.4h | 0.940 | 0.942 | 0.942 |
| WBCEL as ES Crit. | 1 | 4h | 0.912 | 0.878 | 0.877 |
| BCE loss | 5 | 5h | 0.944 | 0.908 | 0.907 |
| WBCE loss with w = 2 | 5 | 5h | 0.944 | 0.914 | 0.913 |
| learining rate 0.001 | 4 | 5.25 | 0.939 | 0.930 | 0.931 |
| use only year $D_{2016}$ | 2 | 2h | 0.914 | 0.927 | 0.926 |

**Table S4:** Model 2, Experimental results

| hyperparameter | epochs | Total time | AUC on Train | AUC on Valid. | AUC on Test |
|---|---|---|---|---|---|
| base | 3 | 6h | 0.92862 | 0.94201 | 0.9428 |
| dropout = 0.5 | 2 | 10h(max) | 0.93471 | 0.94981 | 0.9510 |
| filters sizes: (5,2,5,5),(2,5,5),(32,3,3) | 2 | 12h (max) | 0.93588 | 0.94741 | 0.9456 |
| num of filters: 8,32,8 | 2 | 10h(max) | 0.94540 | 0.95599 | 0.9563 |
| SPP with n=2 and k = $[12, 5]$ | 4 | 10h(max) | 0.94666 | 0.95731 | **0.9569** |
| Cost as ES | 3 | 10h(max) | 0.94196 | 0.93736 | 0.9371 |
| In size 31 | 5 | 10h(max) | 0.94359 | 0.95100 | 0.9509 |
| In size 45 | 3 | 10h(max) | 0.94132 | 0.95189 | 0.9514 |
| $\theta$=0.5 | 4 | 5h | 0.93257 | 0.94898 | 0.9479 |
| wBCE loss, w= 10 | 3 | 10h(max) | 0.93109 | 0.93944 | 0.9393 |
| L2 reg. with 0.6 | 2 | 10h(max) | 0.92067 | 0.92135 | 0.9203 |
| ADAM lr = 0.001 | 4 | 10h(max) | 0.94794 | 0.93456 | 0.9348 |

**Table S5:** Model 3, Experimental results

| hyperparameter | epochs | Total time | AUC on Train | AUC on Valid. | AUC on Test |
|---|---|---|---|---|---|
| base | 3 | 6h | 0.92862 | 0.94201 | 0.9428 |
| dropout = 0.5 | 5 | 10h(max) | 0.93047 | 0.94497 | 0.9457 |
| hidden dim=(8,8,4,4) | 5 | 10h(max) | 0.92581 | 0.94429 | 0.9429 |
| hidden dim=(16,8,4,4) | 3 | 10h(max) | 0.92859 | 0.94707 | 0.9474 |
| hidden dim=(16,16,4,4) | 3 | 8h | 0.92962 | 0.94573 | 0.9464 |
| Cost Matrix as ES | 5 | 5h | 0.93768 | 0.93787 | 0.9378 |
| size = 31 | 4 | 10h(max) | 0.93143 | 0.94165 | 0.9417 |
| size = 35 | 4 | 9h | 0.93248 | 0.94808 | **0.9484** |
| $\theta = 0.5$ | 3 | 2h | 0.90706 | 0.92173 | 0.9361 |
| wBCE loss with w = 10 | 4 | 6h | 0.91791 | 0.94352 | 0.9423 |
| L2 regularization with param 0.6 | 2 | 10h(max) | 0.92067 | 0.92135 | 0.9203 |
| ADAM lr = 0.001 | 4 | 10h(max) | 0.94794 | 0.93456 | 0.934 |

**Table S6:** Model 4, Experimental results

| hyperparameter | epochs | Total time | AUC on Train | AUC on Valid. | AUC on Test |
|---|---|---|---|---|---|
| base | 4 | 5h | 0.92831 | 0.94184 | 0.9420 |
| dropout = 0.5 | 5 | 8h | 0.93149 | 0.94147 | 0.9413 |
| hidden dim= (16,(8,8),8,8) | 2 | 10h(max) | 0.92767 | 0.93447 | 0.9350 |
| hidden dim=(16,(16,16),8,8) | 2 | 10h(max) | 0.92766 | 0.94244 | 0.9433 |
| hidden dim= (8,(8,8),4,4) | 5 | 5h | 0.92678 | 0.94099 | 0.9408 |
| Cost Matrix as ES | 5 | 5h | 0.93565 | 0.94194 | 0.9417 |
| $\theta = 0.5$ | 3 | 3h | 0.91713 | 0.93774 | 0.9363 |
| wBCE loss with w = 10 | 3 | 5h | 0.91369 | 0.92903 | 0.9295 |
| L2 regularization with param 0.6 | 1 | 10h(max) | 0.88856 | 0.91088 | 0.9102 |
| ADAM lr = 0.001 | 5 | 7 | 0.94709 | 0.95324 | **0.9540** |

#### S7.2 Testing on year beyond training period (2018)

Our four best models of each class are compared on labels corresponding to year 2018, $\hat{Y}^j_{2018}, j \in \mathbf{J}_{2017}$. The input data for our last three models, Model 2, Model 3, Model 4, is $(S^j, X^j_{2015}, X^j_{2016}, X^j_{2017)|j\in\mathbf{J}_{2017}}$ whereas Model 1 takes as an input $(S^j, X^j_{2017)|j\in\mathbf{J}_{2017}}$.

**Scenario 1:** Ratio of forested to deforested pixels in 2018 (0:1 labels) is 1:1 and our model predicts the top 50% of the data it receives as deforested. The data size after undersampling with ratio 1:1 is 572978.
**Scenario 2:** Ratio of forested to deforested pixels in 2018 (0:1 labels) is 1:4 and our model predicts the top 20% of the data it receives as deforested. The data size after undersampling with ratio 4:1 1432445
**Scenario 3:** Ratio of forested to deforested pixels in 2018 (0:1 labels) is 9:1 and our model predicts the top 20% of the data it receives as deforested. The data size after undersampling with ratio 9:1 2864890

We ran this test on the three best models, form class Model 1: 2D CNN, Model 2: 3D CNN, and Model 4: Deep ConvLSTM RNN model. Figure S16 , S17 and S18 summarize the our final results. Overall, all model preserve their accuracy when tested on the subsequent year and achieve similar accuracy. Nevertheless, Model 2: 3D CNN achives highest score on Scenario 3, which approximate the actual imbalance of forested to deforested pixels the most.

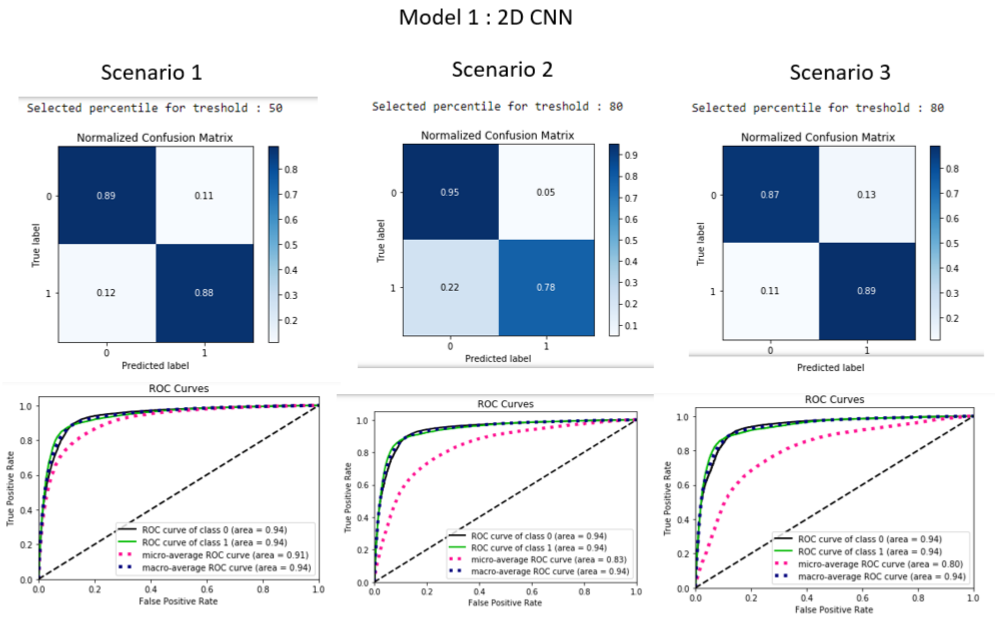

**Figure S16:** Model 1: 2D CNN

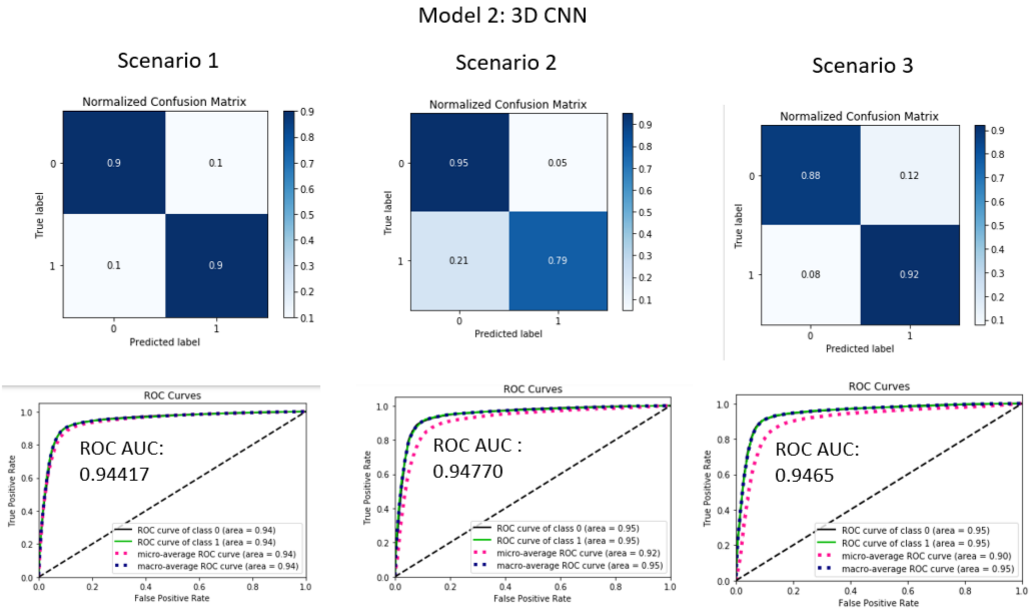

**Figure S17:** Model 2: 3D CNN

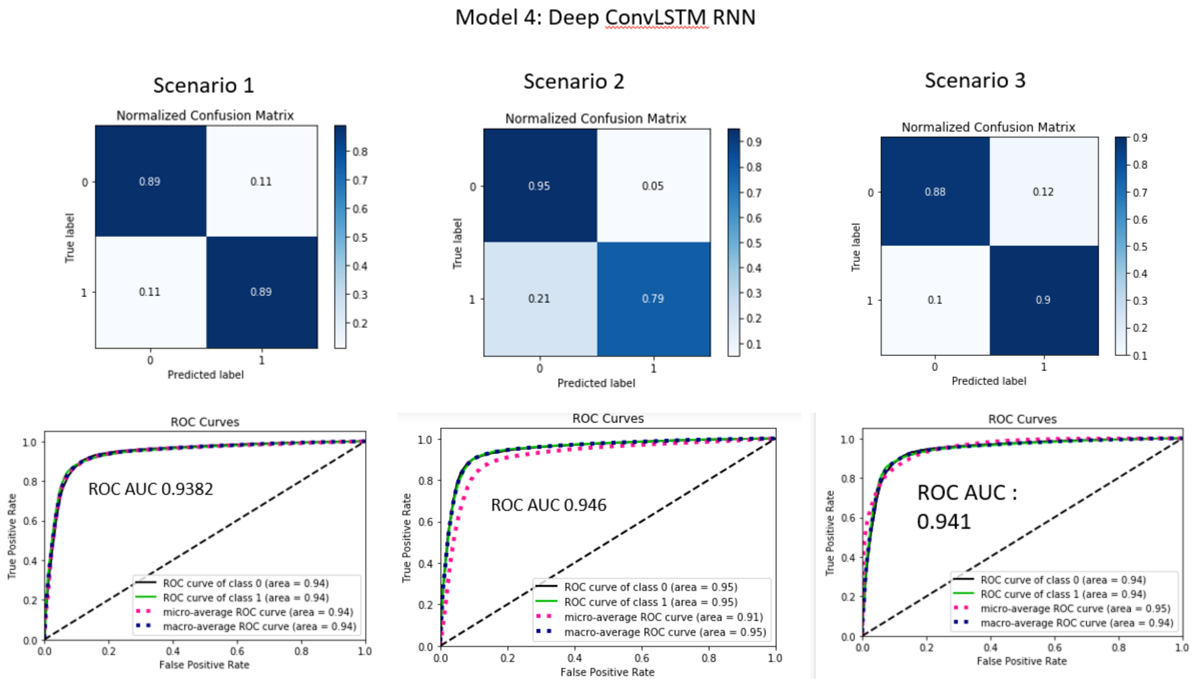

**Figure S18:** Model 4:Deep ConvLSTM RNN

##### S7.2.1 Notes on initial experimentation

The spatial size of input matters in 2D CNNs. For almost all models increasing the number of filters in their layer is beneficial. Setting weighted BCE loss with high weight penalty to wrongly predicted deforested labels, or sampling the model in favour for the deforested pixels ($\theta = 0.5$ - ratio 0:1 labels) does not show any noticeable difference in accuracy. The best performing model is Model 2: 3D CNN, followed by Model 4 and 3. Model 1 is ranked as last.

For our first Model 1, 2D CNN, that the size of the input image is crucial. Secondly, increasing the networks' depth also resulted in better performance on test data. The way we employed dropout did not have noticeable influence, moreover, when it was completely switched off, the accuracy on test and validation data was higher, and the opposite trend was observed when dropout was set to 0.5. Therefore, we can be confident that the way we utilize it is efficiently. Finally, training with smaller learning rate results in a comparable accuracy for the benefit of the faster learning process. We believe this is possible due to the Bach Normalization Layer after each activation.

Our second Model, 3D CNN, noticeably outperformed Model 1 in all experimental scenarios and we confidently conclude that the utilization of the time domain is of huge importance. For this model, the size of the input image is not noted to make huge difference when set to 31, 35 or 45. However, increasing the number of filters in the 3D CNN leads to better accuracy. Furthermore, if the downsampling rate is at a lower rate, performance improves. The wast two facts give us the intuiting that the model has much more information to learn and a deeper network may even result in a higher accuracy.

The third model, ConvLSTM RNN performs almost as well as model 2. All figures about AUC ROC on test data are very similar and clear conclusions can not be made. The only clear conclusion we can make is that including weight decay, also known as L2 penalty resulted in the worst performing model in each model class, although applied with the small parameter 0.6.

Our best model of class Model 4, Deep ConvLSTM model, resulted by a training process when the learinig rate was set to 0.001, and is the second best model overall. The inclusion of additional ConvLSTM cell did not show any significant improvement.

Overall, we conclude that our second proposed model, 3D CNN, achieves the best AUC measurement on the 2017 test data set. It is also simpler then our Model 3 and 4. Furthermore, due to its overall highest performance in all test scenarios, we believe the convergence is not by “chance”.

#### S7.3 Upgraded models: Junin region

##### S7.3.1 2D CNN

Training and testing plots and accuracy statistics are available on WandB [1]

##### S7.3.2 3D CNN

Training and testing plots and accuracy statistics are available on WandB [2]

[1] `https://wandb.ai/patball/forecasting2D`
[2] `https://wandb.ai/patball/forecasting`

### References

*Ballas Nicolas, Yao Li, Pal Chris, Courville Aaron*. Delving Deeper into Convolutional Networks for Learning Video Representations. 2015. pages 19

*Buda Mateusz, Maki Atsuto, Mazurowski Maciej A.* A systematic study of the class imbalance problem in convolutional neural networks // Neural Networks. Oct 2018. 106. 249–259. pages 29

*Fei-Fei Li Silvio Savarese. Juan Carlos Niebles*. STANFORD VISION AND LEARNING LAB CS231n Convolutional Neural Networks for Visual Recognition. 2017. pages 8, 9, 14

*Goldman Elizabeth, Weisse Mikaela*. Global Forest Watch Technical Blog: Global Forest Watch's 2018 Data Update Explained. 2019. pages 3

*Goodfellow Ian, Bengio Yoshua, Courville Aaron*. Deep Learning. 2016. `http://www.deeplearningbook.org`. pages 7, 17

*Hansen M. C., Potapov P. V., Moore R., Hancher M., Turubanova S. A., Tyukavina A., Thau D., Stehman S. V., Goetz S. J., Loveland T. R., Kommareddy A., Egorov A., Chini L., Justice C. O., Townshend J. R. G.* High-Resolution Global Maps of 21st-Century Forest Cover Change // Science. 2013. 342, 6160. 850–853. pages 2, 3, 4, 5

*He Kaiming, Zhang Xiangyu, Ren Shaoqing, Sun Jian*. Spatial Pyramid Pooling in Deep Convolutional Networks for Visual Recognition // Lecture Notes in Computer Science. 2014. 346–361. pages 14, 15

Rectified linear units improve restricted boltzmann machines vinod nair. // . 2010. pages 11

*Hochreiter Sepp, Schmidhuber Jürgen*. Long Short-term Memory // Neural computation. 12 1997. 9. 1735–80. pages 17

*Hoffman Garrett*. Introduction to LSTMs with TensorFlow. 2018. pages 18

*Ioffe Sergey, Szegedy Christian*. Batch normalization: Accelerating deep network training by reducing internal covariate shift // arXiv preprint arXiv:1502.03167. 2015. pages 11, 12

*Kaggle* . 3D Convolutions : Understanding and Implementation. 2018. pages 10

*Kingma Diederik P, Ba Jimmy*. Adam: A method for stochastic optimization // arXiv preprint arXiv:1412.6980. 2014. pages 30, 35

*Li Xiang, Chen Shuo, Hu Xiaolin, Yang Jian*. Understanding the disharmony between dropout and batch normalization by variance shift // arXiv preprint arXiv:1801.05134. 2018. pages 13, 21

*Li Ying, Zhang Haokui, Shen Qiang*. Spectral–Spatial Classification of Hyperspectral Imagery with 3D Convolutional Neural Network // Remote Sensing. 2017. 9, 1. pages 10

*Rußwurm Marc, Körner Marco*. Multi-Temporal Land Cover Classification with Sequential Recurrent Encoders // ISPRS International Journal of Geo-Information. Mar 2018. 7, 4. 129. pages 19

*SHI Xingjian, Chen Zhourong, Wang Hao, Yeung Dit-Yan, Wong Wai-kin, WOO Wang-chun*. Convolutional LSTM Network: A Machine Learning Approach for Precipitation Nowcasting // Advances in Neural Information Processing Systems 28. 2015. 802–810. pages 19

*Snoek Jasper, Larochelle Hugo, Adams Ryan P.* Practical bayesian optimization of machine learning algorithms // Advances in neural information processing systems. 2012. 2951–2959. pages 28

*Srivastava Nitish, Hinton Geoffrey, Krizhevsky Alex, Sutskever Ilya, Salakhutdinov Ruslan*. Dropout: a simple way to prevent neural networks from overfitting // The Journal of Machine Learning Research. 2014. 15, 1. 1929–1958. pages 12
